# Distinguish risk genes functioning at presynaptic or postsynaptic regions and key connectomes associated with pathological α-synuclein spreading

**DOI:** 10.1101/2025.03.11.642462

**Authors:** Yuanxi Li, Justin Torok, Jessica Ding, Ning Wang, Courtney Lau, Shruti Kulkarni, Chaitali Anand, Julie Tran, Michael Cheng, Claire Lo, Binbin Lu, Yanzi Sun, Xia Yang, Ashish Raj, Chao Peng

## Abstract

Previous studies have suggested that pathological α-synuclein (α-Syn) mainly transmits along the neuronal network, but several key questions remain unanswered: (1) How many and which connections in the connectome are necessary for predicting the progression of pathological α-Syn? (2) How to identify risk gene that affects pathology spreading functioning at presynaptic or postsynaptic regions, and are these genes enriched in different cell types? Here, we addressed these key questions with novel mathematical models. Strikingly, the spreading of pathological α-Syn is predominantly determined by the key subnetworks composed of only 2% of the strongest connections in the connectome. We further explored the genes that are responsible for the selective vulnerability of different brain regions to transmission to distinguish the genes that play roles in presynaptic from those in postsynaptic regions. Those risk genes were significantly enriched in microglial cells of presynaptic regions and neurons of postsynaptic regions. Gene regulatory network analyses were then conducted to identify ‘key drivers’ of genes responsible for selective vulnerability and overlapping with Parkinson’s disease risk genes. By identifying and discriminating between key gene mediators of transmission operating at presynaptic and postsynaptic regions, our study has demonstrated for the first time that these are functionally distinct processes.

## Introduction

Parkinson’s disease (PD) is characterized by the accumulation of misfolded alpha-synuclein (α-Syn) in the central nervous system (CNS) (Peng et al., 2018; Peng et al., 2020; Spillantini et al., 1998). Previous evidence suggests that PD progression is driven by the intercellular spreading and templated amplification of pathological α-Syn in the CNS (El-Agnaf et al., 2006; Mollenhauer et al., 2008; Peng et al., 2018). Hence, blocking either the spreading or amplification of pathological α-Syn are promising therapeutic strategies for PD (Peng et al., 2020). Injecting misfolded α-Syn preformed fibrils (PFF, ‘pathological seeds’) into the brains of wild-type (WT) mice can induce misfolding of native α-Syn into pathological conformations, which then spreads to other brain regions (Luk et al., 2012a). This injection mouse model for PD research is ideal for studying the spreading of pathological α-Syn along the neuronal network (Henderson et al., 2019a; Luk et al., 2012a).

Biophysical or mathematical models of network spreading have become useful tools to augment experimental studies. Such models can facilitate our understanding of rich experimental data by helping to validate specific hypotheses about disease mechanisms that are difficult to observe directly. Biophysical models can further reveal which processes control disease dynamics, and uncover key mechanisms that future treatments could be designed to disrupt (Torok et al., 2023). Mathematical models, especially the network diffusion model (NDM) (Raj et al., 2012), the Smoluchowski network model (SNM) (Bertsch et al., 2021; Fornari et al., 2020; Raj et al., 2021), and the Nex*is* model (Anand et al., 2022), which rely upon the mesoscale connectome (Knox et al., 2018; Moon et al., 2014) and Allen Gene Expression Atlas (Lein et al., 2007), have been used *in silico* to model and predict the progression of pathological proteins. Pathological α-Syn progression was modeled based on the following factors: pathology amplification and clearance, network spread, and regional selective vulnerability explained by individual gene expression (Anand et al., 2022; Dadgar-Kiani et al., 2022; Henderson et al., 2019a; Mezias et al., 2020).

Despite the progress made by previous studies (Dadgar-Kiani et al., 2022; Henderson et al., 2019a; Henderson et al., 2020; Mezias et al., 2020; Pandya et al., 2019), the following important questions remain unclear: 1. Previous models are based on the hypothesis that pathological α-Syn spreading is either fully anterograde or fully retrograde in WT mice and *G2019S LRRK2* transgenic mice (Henderson et al., 2019a). However, bidirectional spreading with both retrograde and anterograde components is likely but has not been investigated in previous studies. What are the proportions of the anterograde spreading and retrograde spreading? 2. Is there any difference in pathological α-Syn spreading directionality preference between WT mice and transgenic mice? 3. All previous studies have used the whole or majority of the whole mesoscale connectome to model the spreading of pathological α-Syn. It remains unknown how many connections of the whole connectome are necessary for predicting the pathological α-Syn progression. In other words, are there subsets of key connections in the connectome that predominantly determine the spreading patterns of pathological α-Syn? 4. In previous studies, genes that contribute to the selective vulnerability of pathological α-Syn spreading are identified based on the assumption that they only function in presynaptic regions (the ‘outgoing connections’, see **Equation** 8 in **Methods**). However, it is possible that distinct genes modulate pathological α-Syn spreading in postsynaptic regions specifically (the ‘incoming connections’); or in both the presynaptic and postsynaptic regions. 5. Both neurons and glial cells have been shown to modulate the spreading of pathological proteins (Peng et al., 2020). It is unknown which cell type predominantly encodes the selective vulnerability of pathological α-Syn spreading.

Here, we quantitatively measured the spatiotemporal patterns of pathological α-Syn in PFF injected WT mice at 3- and 6- months-post-injection (MPI) at a higher resolution than the previous study (Henderson et al., 2019a). The data was coupled with the augmented Nex*is* model (Anand et al., 2022) that incorporated the templated amplification, pathology clearance, connectome-based directionality-biased spreading and regional gene expression patterns (See **Equations** 1-7 in **Methods**), and systems biology approach (See **Systems biology analyses** in **Methods**) to address the questions above. Using our new mathematical model and datasets from a previous study (Henderson et al., 2019a), we found that pathological α-Syn transmitted predominantly retrogradely in WT mice, but the directionality of pathological α-Syn spreading in *G2019S LRRK2* mice was bidirectional with a slight anterograde bias. This is the first report on the quantitative proportion of pathological α-Syn spreading and how *G2019S LRRK2* affects pathological α-syn spreading directionality, giving a more nuanced picture of the directional preference of α-Syn than previously assumed (Dadgar-Kiani et al., 2022; Henderson et al., 2019a; Mezias et al., 2020; Pandya et al., 2019). Unexpectedly, we then found that the spread of pathological α-Syn can be predicted using as few as the top 2% strongest connections of the whole mesoscale connectome. This surprising finding suggests that the spreading of pathological α-Syn is predominantly determined by key connectomes composed of a minor subset of brain regions and connections, which will be very useful evidence for experimental scientists to focus their research on these ‘key connectomes’. We also identified candidate risk genes that modulated pathological α-Syn spreading by affecting the presynaptic regions specifically, postsynaptic regions specifically, and both (See **Equation** 8 in **Methods**). We performed systems biology analyses to study the similarities and differences of top risk genes ranked by our model performance. We found that the risk genes of outgoing effect were enriched in microglial cells, while the risk genes of incoming effect were enriched in neurons, indicating that different cell types play distinct roles in determining the selective vulnerability at presynaptic versus postsynaptic sites. Gene regulatory network analyses were then conducted to identify ‘key drivers’ of genes responsible for selective vulnerability and overlapping with Parkinson’s disease risk genes. Overall, our study successfully investigated the key connectomes and risk genes at different synaptic sites in modulating α-Syn pathology progression. From a computational aspect, our study provided a pipeline to predict and distinguish the vulnerable genes from presynaptic or postsynaptic regions of PD, and inspired research on other neurodegenerative diseases.

## Results

### Quantitative pathology in the stereotaxic injection mouse model

We used the stereotaxic injection mouse model to study the progression of pathological α-Syn in the CNS (Luk et al., 2012a). Pathological α-Syn PFF was stereotaxically injected into the ipsilateral dorsal striatum (ipsilateral caudoputamen region, iCP) of two-to-three-month-old C57BL6/C3H wild-type mice to mimic the progression of α-Syn pathology. Mice were euthanized at 3 and 6 months-post-injection (MPI) for the immunohistochemistry analyses (**Fig. 1A**). α-Syn pathology was revealed by staining with antibody against pathological α-Syn (Syn506) (Peng et al., 2018). Stained brain slices at 9 specific bregma levels were selected per mouse and annotated according to the Allen Mouse Brain Reference Atlas (AMBA, https://mouse.brain-map.org/static/atlas). The QuPath software was then used to quantify the percentage area occupied by α-Syn pathology in 196 brain grey matter regions (**Fig. 1B**). With the updated AMBA database, we were able to achieve a higher resolution (Henderson et al., 2019a).

**Figure 1.**
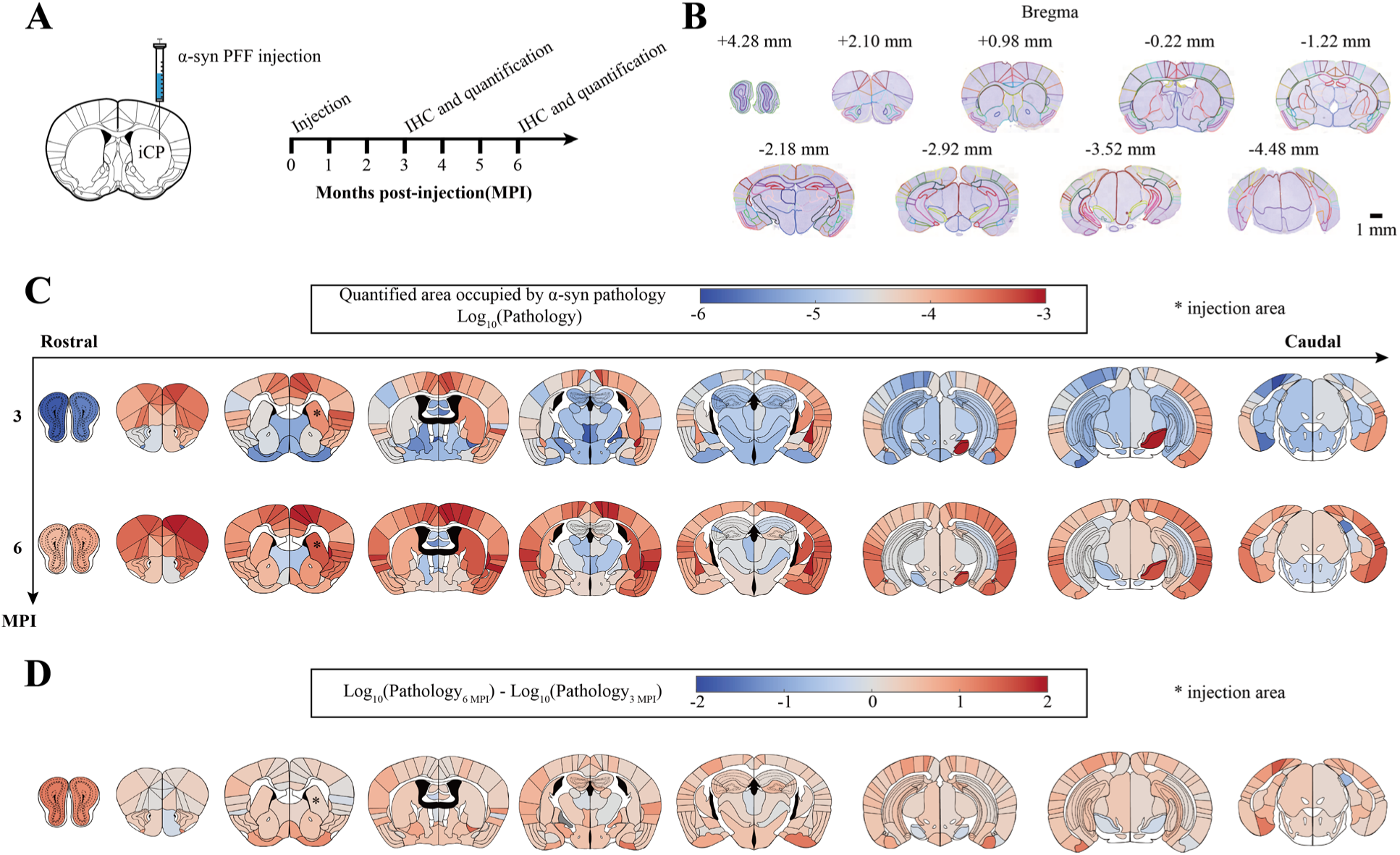
Immunohistochemistry experiments and quantitative biology. **A**, Experiment schematic: pathological α-Syn PFF was stereotaxically injected into the iCP region. Mice were euthanized at 3 and 6 months-post-injection prior to immunohistochemistry, which were then quantitatively assessed for α-Syn inclusions using QuPath. **B**, Representative images of brain sections and annotations. 9 brain bregma level were selected per mouse, and 196 grey matter regions were manually annotated. **C**, Heatmap of regions affected by α-Syn pathologies (log10-transformed; *, injection area). Warmer colors represent areas with higher pathology burdens, while cooler colors represent areas with lower pathology burdens. **D**, Differences of regional pathology burdens between 3 and 6 MPI, where the color indicates the fold change in regional pathology burden between 3 and 6 MPI. Scale bar in **B**: 1 mm.

The spatiotemporal distributions of α-Syn pathology are shown in **Fig. 1C** (See also **Supplementary Table - Quantitative Pathology**). Similar to previous findings (Henderson et al., 2019a; Peng et al., 2018), the ipsilateral side had more pathology than the contralateral side (**Fig. 1C**). Comparisons between 3 MPI and 6 MPI (**Fig. 1D**) showed distinct pathological burden changes in subregions of hippocampus, prefrontal cortex, and midbrain. As reported before, decreased pathology burden was observed in the substantia nigra (SN) between 3 and 6 MPI (Henderson et al., 2019b; Luk et al., 2012a). With our higher annotation resolution, we also found reduced pathology in the ipsilateral postsubiculum (iPOST) (**Fig. 1D**, **Fig. S1**), but vast majority of regions showed increased α-Syn burdens between 3 MPI and 6 MPI (**Fig. 1D**). We also identified a significant increase in pathology burden in the ipsilateral olfactory areas (iOLF), the primary motor area (iMOp), the ipsilateral primary somatosensory area, lower limb (iSSp-ll), and the contralateral caudoputamen (cCP) (**Fig. S1-S3**, **Supplementary Table - Quantitative Pathology**).

### Different directionality preferences of pathological α-Syn spreading in WT mice and *G2019S LRRK2* transgenic mice

A very important question regarding pathology spread is the preference of direction (anterograde vs. retrograde spread). Previous mathematical models of pathological α-Syn spreading have only compared the situation of fully anterograde transport and fully retrograde transport, and concluded that pathological α-Syn spreading is due to fully retrograde transport in both WT mice and *G2019S LRRK2* mice (See **Equation** 4 in **Methods**) (Dadgar-Kiani et al., 2022; Henderson et al., 2019a; Mezias et al., 2020); however, while both anterograde and retrograde transport have been reported for pathological α-Syn (Freundt et al., 2012), unequal bidirectional spread has not been previously explored in mathematical models (Dadgar-Kiani et al., 2022; Henderson et al., 2019a; Mezias et al., 2020; Pandya et al., 2019).

We used Nex*is*:global (Anand et al., 2022) as the global spread model to investigate the pathological progression including the amplification, clearance, and spreading of pathological α-Syn (See **Equation** 1- 7 in **Methods** for details). To accommodate a continuous measure of net directional preference, the original Nex*is*:global model (Anand et al., 2022) was augmented by introducing a new parameter, *s*, to indicate spreading direction. *s* is bounded between 0 and 1, with 0 indicating fully anterograde spread and 1 indicating fully retrograde spread (See **Equation** 1-7 in **Methods** for details). To the best of our knowledge, our model has incorporated and modeled the spread directionality preference as a factor in the progression of pathological α-Syn for the first time. We used concordance correlation coefficients averaged over 3 and 6 MPI (average CCC, abbreviated as Ave. CCC in the following paragraphs) as the loss function to fit our model because of it can capture the actual values of the data as well as the trend (scale) (Lawrence and Lin, 1989).

Our new model demonstrates that pathological α-Syn mainly spreads retrogradely with minor anterograde spreading ( *s* = 0.87, Ave. CCC = 0.622) in WT mice, and the overall balance between amplification and clearance was weighted towards amplification, indicated by its positive sign ( *α* = 0.33, see **Methods**) (**Fig. 2A**, **Fig. S4**). We then tested our model under three directionality cases: fully anterograde, fully retrograde, and equal amount of anterograde versus retrograde spread (unbiased spread) (**Fig. 2, B-D**), which all had lower Ave. CCC values than the best directionality condition. To compare with previous methods utilizing Pearson’s correlation coefficient (Pearson’s R) as the target metric (Henderson et al., 2019a; Mezias et al., 2020; Raj et al., 2012; Raj et al., 2015), we found that the average Pearson’s R for our model was approximately the same as Ave. CCC, indicating that our model captures the scale of the observed data (**Fig. 2A**). Using Pearson’s R instead of CCC for parameter fitting destroyed the model’s ability to predict both trend and scale (**Fig. S5**). Our model also accurately captured the disproportionate pathological α-Syn deposition in the ipsilateral hemisphere relative to the contralateral hemisphere at both 3 and 6 MPI (**Fig. S6**).

**Figure 2.**
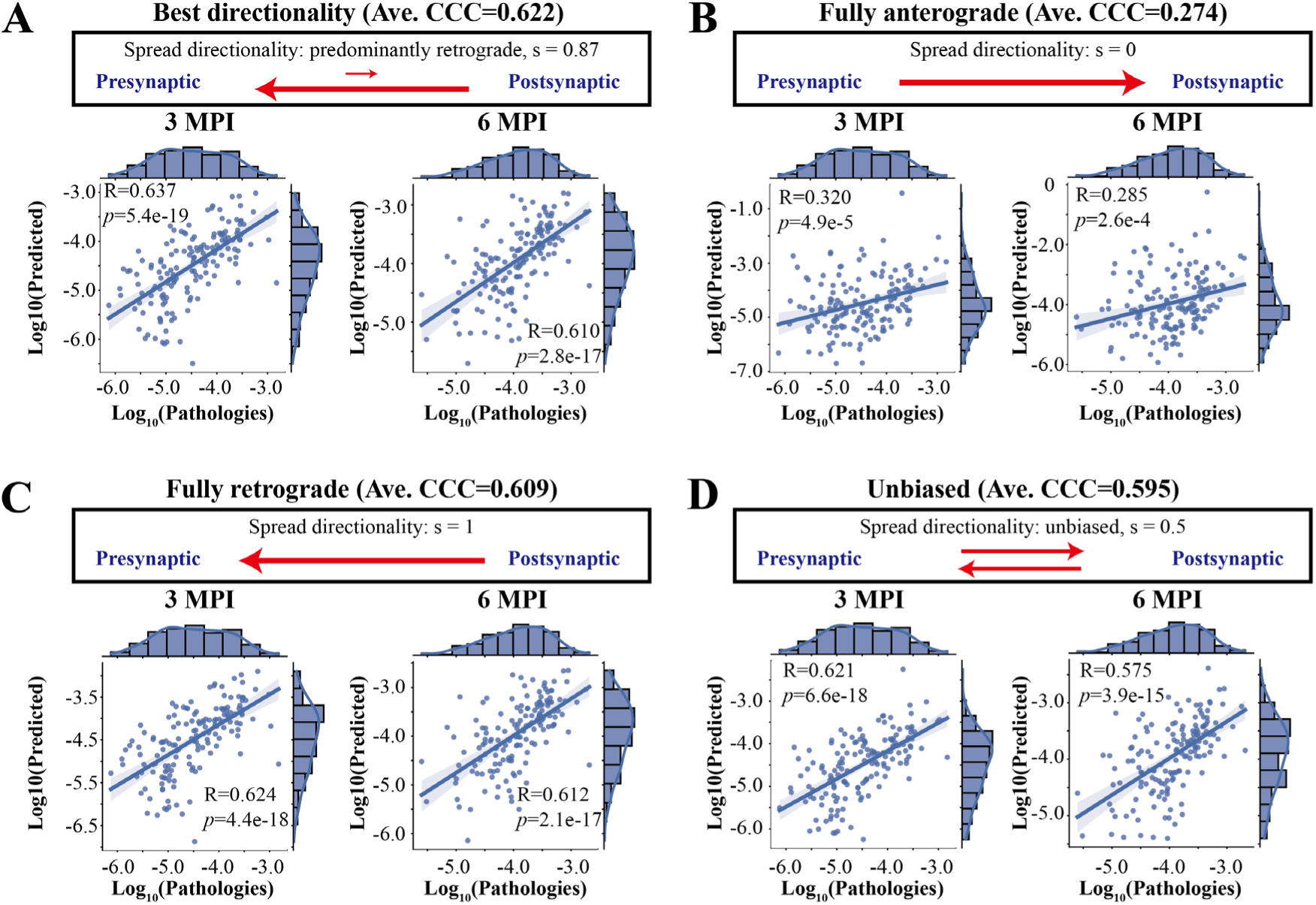
Global spread model based on the mesoscale connectome revealed the spread directionality of α-Syn was predominantly retrograde with a minor anterograde component. **A**, Best model results (Ave. CCC = 0.622) with the directionality parameter of s=0.87. **B**, Model results (Ave. CCC = 0.274) with the fully anterograde parameter of s=0. **C**, Model results (Ave. CCC = 0.609) with the fully retrograde parameter of s=1. **D**, Model results (Ave. CCC = 0.595) with the unbiased directionality parameter of s=0.5. The directionality of pathological α-Syn spread is shown by the arrows in the box, while the length of the line indicates the proportion of spread direction. Each dot represents one brain region and the x-axis and y-axis represent the pathology (log10-transformed) found empirically and predicted by the model, respectively. For each situation, the Pearson’s correlation coefficient and the best regression lines for both 3 and 6 MPI are also displayed. The shaded ribbon represented the 95% prediction interval. Abbreviations: R, Pearson’s correlation coefficient; *p*, *p* values from linear regression.

To validate our observation that pathological α-Syn mainly spread retrogradely in WT mice, we then tested the spread directionality using our model and previous datasets from Henderson et al. (Henderson et al., 2019a), which consisted of two groups of mice (wild-type nontransgenic mice and *G2019S LRRK2* mice; see **Methods** for further details). In agreement with the results above, our model predicted that pathological α-Syn also spread predominantly retrogradely in these WT mice (**Fig. S7A**, Ave. CCC=0.599, *s* = 0.75). More interestingly, we found that the directionality of pathological α-Syn spread in *G2019S LRRK2* transgenic (*G2019S*) mice was very different from that in the WT mice, showing bidirectional spread with a slight anterograde preference (**Fig. S7B**, Ave. CCC=0.561, *s* = 0.40). We also tested the previous hypothesis of retrograde spread (Henderson et al., 2019a) in *G2019S LRRK2* mice (**Fig. S7C**, Ave. CCC=0.508, *s* = 1), which performed worse than the best directionality model (**Fig. S7B**). We conclude that the *G2019S LRRK2* modification has a dramatic effect on the directionality of pathological α-Syn spread.

### The spread of pathological α-Syn is predicted using only 2% connections of the whole connectome

To evaluate whether the actual connectome best predicts the spreading of pathological α-Syn, we tested if random artificial connectomes (‘null models’) could predict pathological α-Syn progression. We fit our data using connectomes with elements sampled from a uniform random distribution, with 1,000 replicates to demonstrate the robustness. As expected, the actual connectome predicted pathological α-Syn spreading much better than all randomly generated matrices (**Fig. 3A**). Similarly, we tested the case when elements of the whole connectome were randomly permutated; the actual connectome also outperformed the permutated connectomes (**Fig. 3B**). These results demonstrate that pathological α-Syn spreads along the actual mesoscale connectome.

**Figure 3.**
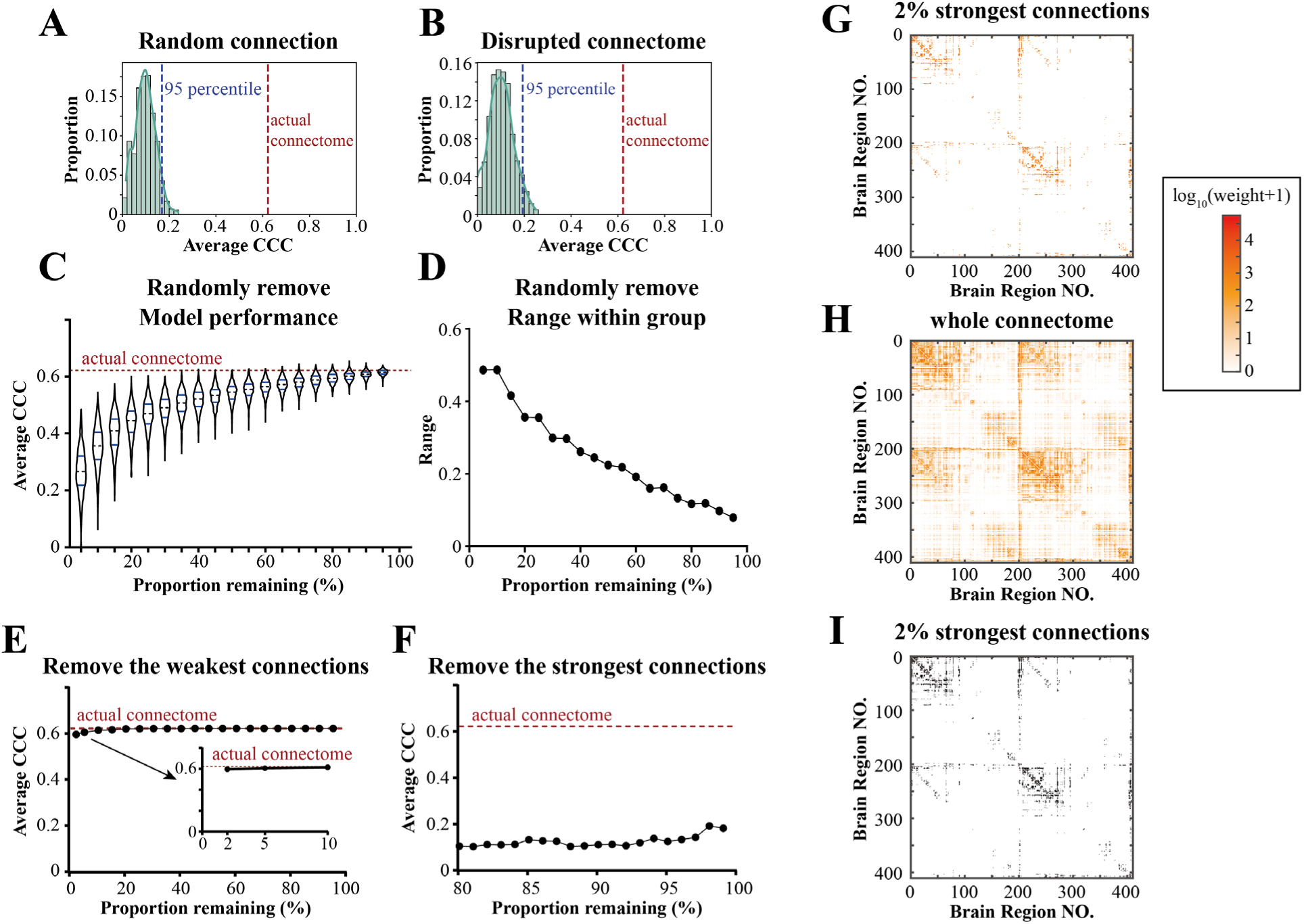
The spread of pathological α-Syn was driven by the partial connectome composed of only top 2% strongest connections. **A**, Distribution of model fitting results using 1,000 matrices with entries sampled from the standard uniform distribution in place of the actual connectome (Ave. CCC at 95th percentile = 0.169). **B**, Distribution of model fitting results using matrices with randomly permuted elements of the actual connectome (Ave. CCC at 95th percentile = 0.194). **C-D**, Distributions of model fitting results by removing random subsets of connections of various sizes at a 5% incremental gradient, resampling the connectome 1,000 times per subset size. **C**, Showing the model performance: For each, the dash line represented the median, and the two solid line represented the 25% and 75% percentiles. **D**, Showing the range (maximum minus minimum Ave. CCC value) within the group. **E-F**, Model fitting results after removing proportions of edges from the connectome in order of connection strength. **E**, Model fitting results after removing from weakest connections; note that the model did not fit successfully when removing 99% of the weakest connections. **F**, Model fitting results after removing the strongest connections. The dotted line of the actual connectome showed the best model fitting of the global spread model with the actual (whole) connectome (Ave. CCC = 0.622). **G-I**, Visualization of key connectomes (2% of the strongest connections), showing heatmaps of the connection values, where the warmer colors represent stronger connections (log10(raw_value+1)-transformed). **G**, Showing 2% of the strongest connections of the connectome. **H**, Showing the whole connectome. **I**, Heatmap of the adjacency matrix corresponding to **G**. The actual connectome here consists of 410 brain regions with 168,100 connections, and 2% of connectome equals to 3,362 connections.

Previous studies have used the whole or a majority of the whole mesoscale connectome to model the spreading of pathological α-Syn (Dadgar-Kiani et al., 2022; Henderson et al., 2019a; Henderson et al., 2020; Mezias et al., 2020; Pandya et al., 2019). However, it remains unknown whether the connections in the connectome contribute equally to the spreading of pathological α-Syn, or if the spreading of pathological α-Syn is predominantly determined by a subset of the connections. To test these hypotheses, we randomly removed different proportions (5%∼95%) of the actual connectome and evaluated how well the remaining partial connectome can predict pathological α-Syn spreading, repeating each case 1,000 times for robustness (**Fig. 3C**). Interestingly, the model worked reasonably well when removing 50%∼60% of the connections, and could achieve great accuracy in some specific cases even with only 5% of connections remaining (**Fig. 3C**). We also observed much higher variation of the performance of the model when more connections were removed (**Fig. 3D**). These results suggest that connections in the connectome may not contribute equally. In other words, there should be subnetworks composed of key connections in the connectome which predominantly determine the spreading pattern of pathological α- Syn.

We hypothesize that the key subnetworks that determine the spreading of pathological α-Syn are composed of the strongest connections in the connectome. To test this hypothesis, we removed different proportions of the strongest or weakest connections from the connectome and evaluated the effects on our model’s performance (**Fig. 3E**, **Fig. 3F**). Strikingly, using as few as 2% of the strongest connections from the actual connectome (3,362 of 168,100 connections), our model still predicted the spreading of pathological α-Syn almost as well as using the complete connectome (**Fig. 3E**). By contrast, removing only the top 1% of the strongest connections from the connectome dramatically disrupted the performance of the model (**Fig. 3F**). Accordingly, we visualized these connections as well as the whole connectome (**Fig. 3, G-I; Fig. S8**).

We then validated this observation using the datasets of WT mice and *G2019S LRRK2* mice (**Fig. S9**) from Henderson et al. (Henderson et al., 2019a). As expected, the progression of pathological α-Syn was well predicted by the 8% of the strongest connections from the ‘masked’ connectome (1,077 of 13,456 connections. **Fig. S9A, Fig. S9C**), and removing even the top 1% of the strongest connections greatly decreased model performance (**Fig. S9B, Fig. S9D**). Note that we used 8% of the strongest connections from the ‘masked’ connectome instead of 2%, because the ‘masked’ connectome used in Henderson et al. (Henderson et al., 2019a) has approximately 13-fold fewer connections than we used in our study. These findings strongly support our hypothesis that the spreading of pathological α-Syn is predominantly determined by a small subset of the strongest connections in the brain (**Fig. 3, Fig. S9**).

### Distinct genes modulate pathological α-Syn spreading on the presynaptic versus postsynaptic site

The spreading pattern of pathological α-Syn has not been fully predicted by the anatomic connectome as well as the global amplification and clearance effects (**Fig. 2A**). This observation suggests that certain brain regions are selectively vulnerable to pathological α-Syn accumulation and spreading, which could be caused by the distinct regional gene expression levels (Alegre-Abarrategui et al., 2019; Henderson et al., 2019a; Peng et al., 2018; Taguchi et al., 2019). To investigate if pathology burden distribution is directly correlated with gene expression levels, gene expression data of 3,855 genes from the AGEA coronal series (Lein et al., 2007) were analyzed, and we measured the gene correlations by calculating Pearson’s correlation coefficient between individual gene expression vector and pathology burden levels of our WT mice. The results showed that at both 3 and 6 MPI, the global spread model was significantly better than the gene correlation model (**Fig. S10**), demonstrating that mesoscale connectome is necessary in modulating pathological α-Syn spreading.

We sought to incorporate gene expression in our global spread model to study whether connectome and gene expression together contribute to α-Syn spread. Previous studies to identify genes that contribute to selective vulnerability have only focused on genes that modulate spreading on the outgoing connections at presynaptic site (See **Equation** 8 in **Methods**) (Dadgar-Kiani et al., 2022; Henderson et al., 2019a; Henderson et al., 2020). However, the effects of gene expression on the postsynaptic site, and on both the presynaptic and postsynaptic regions, have not been previously considered. To identify genes that affect the outgoing connectome alone (i.e., the ‘outgoing’ effect, which models the effect of gene expression on the presynaptic regions alone), incoming connections alone (i.e., the ‘incoming’ effect, which models the effect of gene expression on the postsynaptic regions alone), or equally on outgoing and incoming connections (i.e., the ‘combined’ effect), we revised and augmented the Nex*is* model (Anand et al., 2022). Briefly, the original connectome matrix incorporated in the model was the same as the global spread model (**Fig. 4A**), and we modified the connectome matrix to incorporate the effect of gene expression on outgoing connections (**Fig. 4B**), incoming connections (**Fig. 4C**), or both (combined effect; **Fig. 4D**) (See **Equation** 8-9 in **Methods**).

**Figure 4.**
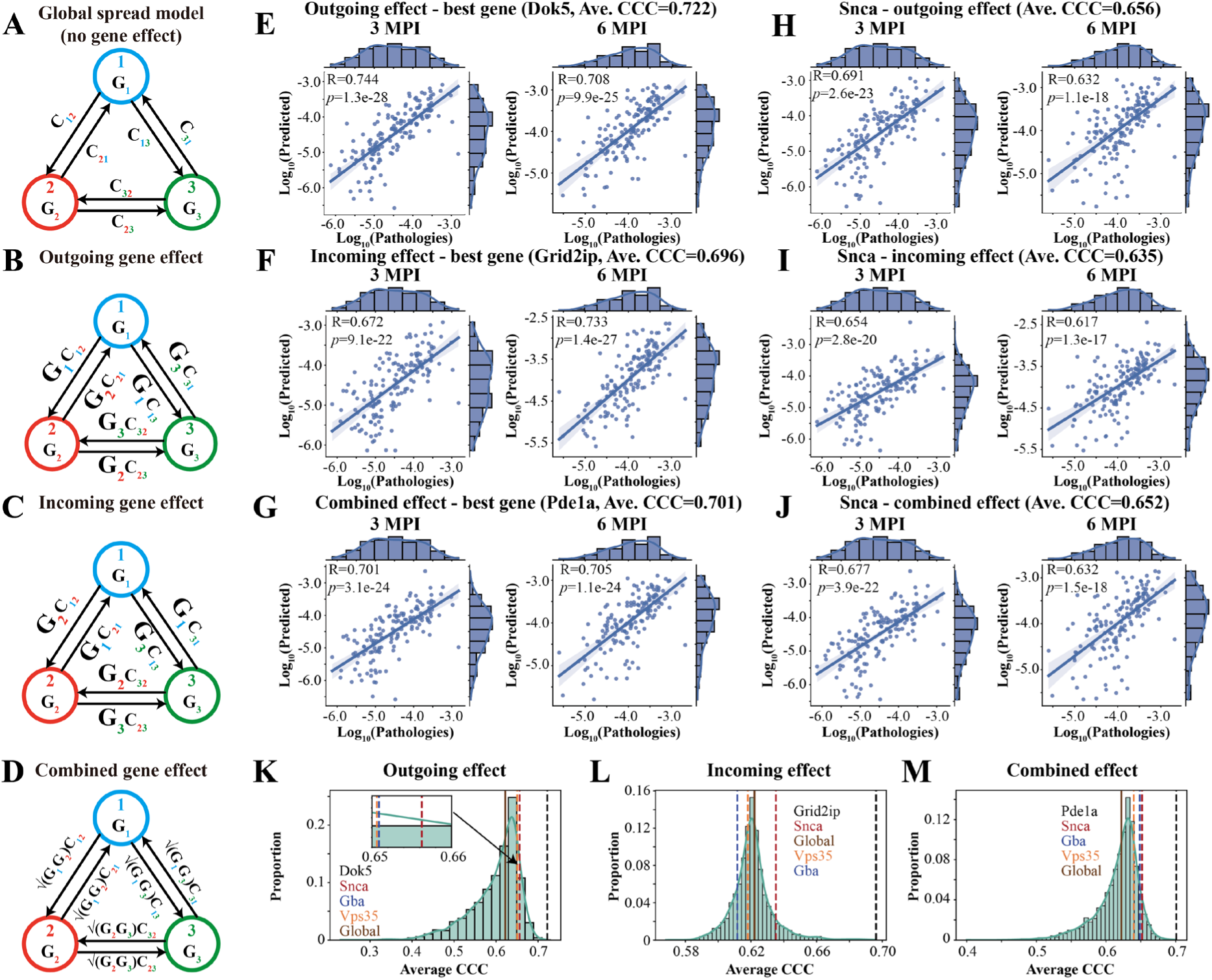
Model results of gene expression model for the outgoing, incoming, and combined effects. **A-D**, Schematics of the global spread model (**A**) and gene expression models (**B**, outgoing effect; **C**, incoming effect; **D**, combined effect): The circles in different colors indicates brain regions; G^i^ in the circles indicate the regional gene expression; and C_ij_ indicate the axonal projection strength from brain region i to j. The regional gene expression had different updated effects on the connectome by revising nothing (**A**, global spread model), revising only the outgoing connections (**B**, outgoing effect), only the incoming connections (**C**, incoming effect), or both equally (**D**, combined effect). **E**, Model results of the best outgoing effect gene (*Dok5*). **F**, Model results of the best incoming effect gene (*Grid2ip*). **G**, Model results of the best combined effect gene (*Pde1a*). **H-J**, Model results of *Snca* gene for its outgoing (**H**), incoming (**I**), and combined effects (**J**), respectively. **K**, The distribution of model results of outgoing effect (model performance: *Dok5* > *Snca* > *Gba* > *Vps35* > *Global*). **L**, The distribution of model results of incoming effect (model performance: *Grid2ip* > *Snca* > *Global* > *Vps35* > *Gba* > *Vps35*). **M**, The distribution of model results of combined effect (model performance: *Pde1a* > *Snca* > *Gba* > *Vps35* > *Global*). Each dot in **E-J** represents one brain region, while the x-axis and y-axis represent the pathology (log10-transformed) found empirically and predicted by the model, respectively. The Pearson’s correlation coefficient and the best regression lines for both 3 and 6 MPI are also displayed. The shaded ribbon in **E-J** represents the 95% prediction interval. Abbreviations: R, Pearson’s correlation coefficient; *p*, *p* values from linear regression.

Based on our WT mice data, individual genes were incorporated in the model, and ranked according to the model performance (**Fig. 4, E-J**; Also see **Supplementary Table - Gene Results)**. We compared the results of these connectome plus gene models with global spread model or gene correlations, showing that our connectome plus gene models predicted the pathological α-Syn spreading better than the other two models (**Fig. S11**). Then, genes that improved the Ave. CCC values over the global spread model were considered to contribute to selective vulnerability of pathological α-Syn spreading. We identified *Dok5* (Ave. CCC=0.722), *Grid2ip* (Ave. CCC=0.696), and *Pde1a* (Ave. CCC=0.701) as the single-best genes for the outgoing, incoming, and combined effect, respectively (**Fig. 4, E-G)**. We also showed the distribution of Ave. CCC values for all genes (**Fig. 4, K-M**). Overall, distinct genes were identified for outgoing, incoming, and combined effects, which suggests that different components of spreading process may be modulated by distinct sets of genes. The top incoming gene, *Grid2ip*, did not contribute to selective vulnerability in the outgoing (ranking 3736^th)^ and combined (ranking 3791^st^) effects. In contrast, *Dok5* and *Pde1a* performed very well for outgoing effect (*Dok5*, ranking 1^st^; *Pde1a*, ranking 53^rd^), incoming effect (*Dok5*, ranking 53^rd^; *Pde1a*, ranking 58^th^), and combined effect (*Dok5*, ranking 2^nd^; *Pde1a*, ranking 1^st^) (See **Supplementary Table - Gene Results**). These results suggest that *Dok5* and *Pde1a* influence both the presynaptic and postsynaptic regions to modulate the spread, whereas *Grid2ip* only influences the postsynaptic regions. Furthermore, similar to the global spread model, the gene model also captured the scale features of the data (**Fig. S12**) (Lawrence and Lin, 1989) and the hemispherical asymmetry of the pathological distribution (**Fig. S13**). We then visualized pathological distribution predicted by these genes (**Fig. S14**), which showed similar trends as the results of the global spread model in the annotated regions (**Fig. S4**). Interestingly, the outgoing and combined effect altered the Ave. CCC values much more than the incoming effect (**Fig. 4, K-M**). The incorporation of the best genes into the model only increased the Ave. CCC values by less than 0.1 (**Fig. 4, E-G**), suggesting that the connectome played a predominant role in the spread of pathological α-Syn.

We also investigated the effects of four important PD genetic risk factors (*Snca*, **Fig. 4, H-J**; *Gba*, **Fig. S15**; *Vps35*, **Fig. S16**; *Gpnmb*, **Fig. S17**). *Snca* modulated both the outgoing (ranking 353^rd^; **Fig. 4H**, **Fig. 4K**) and incoming effects (ranking 356^th^; **Fig. 4I**, **Fig. 4L**), supporting that the expression of *Snca* affected not only the presynaptic but also the postsynaptic sites. Also, *Snca* was ranked higher for all the three effects than *Gba* and *Vps35* (**Fig. 4, K-M)**, demonstrating greater importance of *Snca* in modulating the spread of pathological α-Syn over *Gba* and *Vps35*. We instead found that *Gba* and *Vps35* mainly modulated the outgoing effect (*Gba*, ranking 497^th^, **Fig. S15A**, **Fig. 4K**; Vps35, ranking 506^th^, **Fig. S16A**, **Fig. 4K**) rather than the incoming effect (*Gba*, ranking 3181^st^, **Fig. S15B**, **Fig. 4L**; *Vps35*, ranking 2370^th^, **Fig. S16B**, **Fig. 4L**), indicating that the *Gba* and *Vps35* genes mainly encoded the selective vulnerability of presynaptic site by altering the regional outgoing connections. Additionally, *Gpnmb* has been reported to participate in the cellular uptake of pathological α-Syn (Diaz-Ortiz et al., 2022). We found that *Gpnmb* improved performance of the outgoing-effect (Ave. CCC=0.637), incoming-effect (Ave. CCC=0.623), and combined-effects (Ave. CCC=0.635) Nex*is* models relative to the global model, validating its importance in the spread of pathology. Compared to the previous studies (Dadgar-Kiani et al., 2022; Henderson et al., 2019a; Henderson et al., 2020), our study provided critical insights into how genetic risk factors affected spreading by identifying effect regions that were presynaptic, postsynaptic, or both.

Further, we sought to investigate whether gene expression altered the spread directionality preference. The probability density of the directionality showed that the spread was still predominantly retrograde after considering the gene effect under all three conditions (**Fig. 5**), confirming our hypothesis that the spread of pathological α-Syn was predominantly retrograde.

**Figure 5.**
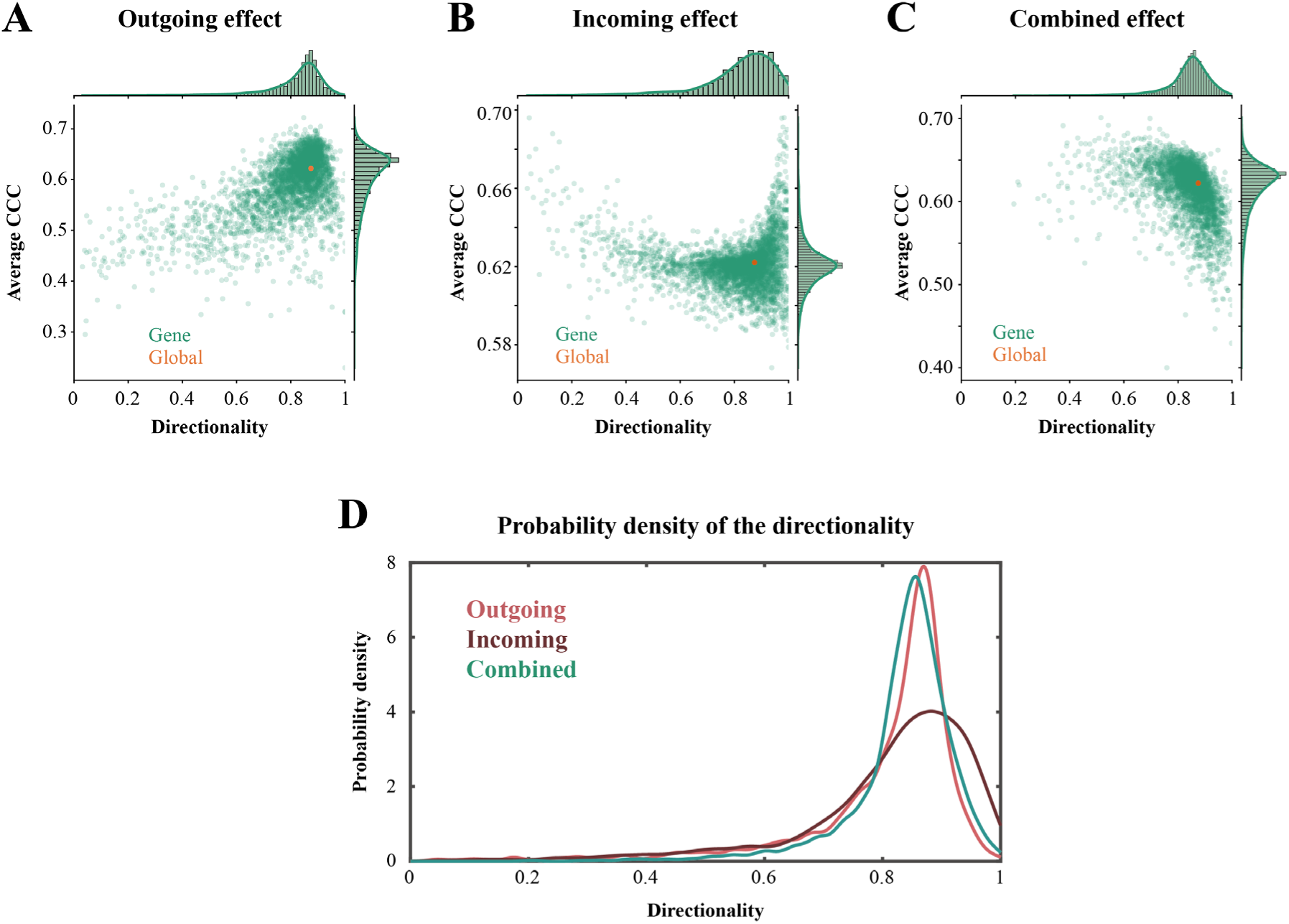
The relationship between individual gene expression and the spread directionality of the gene model. **A-C**, Directionality vs. Ave. CCC, where each green dot represents one gene and the orange dot represents the best results for the global spread model. The x-axis and y-axis represent their directionality parameter values and Ave. CCC, respectively. **D**, Probability density of the directionality of the three groups, where the x-axis and y-axis represent the directionality parameter values and the probability density, respectively.

To study the similarities and differences between the subsets of genes contributing to selective vulnerability of outgoing, incoming, and combined effects, we selected the top 500 genes of the three conditions as the candidate risk genes for further investigation (**Fig. 6A**). In order to ensure that these genes all had robust performance in predicting the spread of pathological α-Syn, we conducted bootstrap analyses for each group’s 500^th^ gene (500th-ranked-gene for the outgoing, incoming, and combined effects were *Large*, *Socs6*, and *Tbc1d14*, respectively) by randomly permuting the elements of the gene expression vectors for 1,000 times. As expected, the performance of the 500^th^ gene’s actual expression values exceeded the 95% percentiles of the permutations for each group (**Fig. S18**), demonstrating that these genes significantly improve model performance relative to a ‘null’ set of genes. We assessed the overlapping of the top 500 genes for the three conditions (**Fig. S19**), which showed a highest overlap between outgoing and combined effects. We also found those genes that strongly modulated the outgoing and incoming effects also modulated the combined effects (**Fig. S19**).

**Figure 6.**
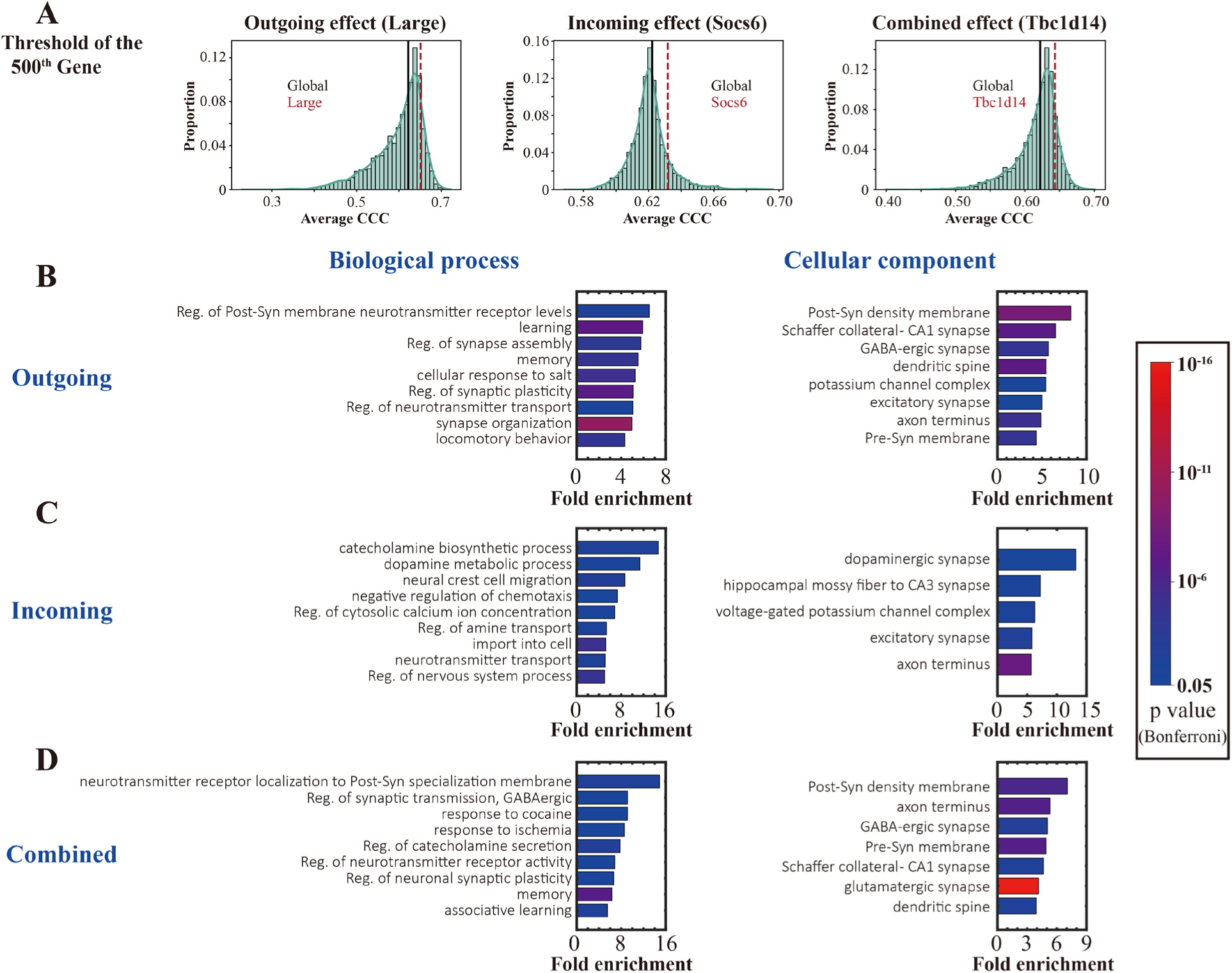
GO analyses for the top 500 genes of outgoing, incoming, and combined effect. **A**, Threshold of the 500th gene of the three groups (*Large* for the outgoing effect, *Socs6* for the incoming effect, and *Tbc1d14* for the combined effect). Each of the three groups exhibited better performance than the global spread model (*p*<0.0001, one sample Wilcoxon test against the Ave. CCC value of global spread model). **B-D**, Biological process and cellular component of GO analyses for the candidate genes from **B**, outgoing effect; **C**, incoming effect; **D**, combined effect. For each GO entry, the displayed results were strictly sorted by the highest fold enrichment and statistically significant after Bonferroni correction. The colors of the bars represent the *p* values, and the length represents the fold enrichment. Only the primary hierarchy is shown. Abbreviations: Reg., regulation; Post-Syn, postsynaptic; Pre-Syn, presynaptic.

We then carried out gene ontology (GO) analyses (Mi et al., 2019) for the top 500 candidate genes (**Fig. 6, B-D; Fig. S20**). Interesting, the candidate genes were all enriched in synaptic components and ion channels such as the dopaminergic (**Fig. 6C**), glutamatergic (**Fig. 6, B-D**), and GABAergic synapses (**Fig. 6B**, **Fig. 6D**), and voltage-gated ion channels (**Fig. S20**). These findings were consistent with the observation that neuronal firing patterns can affect the spread of pathological α-Syn (Kulkarni et al., 2022; Wu et al., 2020; Yamada and Iwatsubo, 2018). Outgoing genes were enriched in biological processes including learning, memory, and locomotory behavior (**Fig. 6B**). By contrast, incoming genes were mainly related to dopaminergic system and synapses (**Fig. 6C**). We therefore find that while there is significant overlap in the functional enrichment of these gene subsets, there are also important differences that suggest that gene modulation of outgoing vs. incoming genes may involve different biological and molecular processes.

### Distinct cell type enrichment for outgoing, incoming, and combined genes

It is unclear which cell types contribute most to the selective vulnerability of pathological α-Syn spreading. To address this question, cell type enrichment analyses (Langlieb et al., 2023) were performed on the top 500 genes for each of the three different effect conditions (**Fig. 7A**, **Supplementary Table - Cell Type Analysis**), which revealed distinct cell type enrichment preference for the different effects. Outgoing genes were significantly enriched in microglia cells, while incoming genes were significantly enriched in neurons (including excitatory and inhibitory neurons). This result suggests that microglial cells at presynaptic sites and neurons at postsynaptic sites show stronger effects in modulating pathological α-Syn spread.

**Figure 7.**
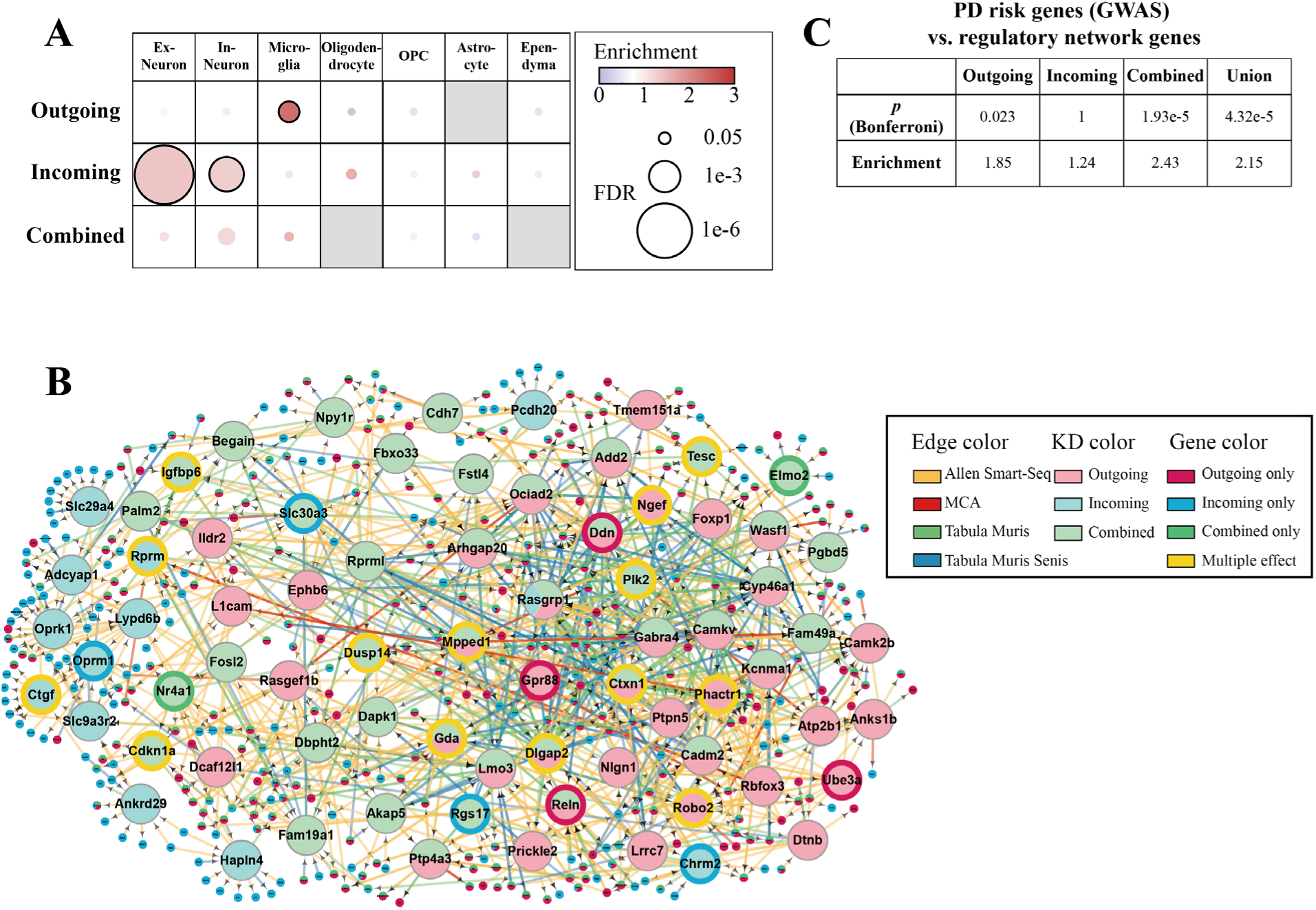
Results of cell-type, key driver (KD), and GWAS-associated enrichment analyses for the candidate (top 500) genes of outgoing, incoming, and combined effects. **A**, Cell-type analyses based on the enrichment of cell type marker genes among the candidate genes of each effect. The sizes of the circles indicate FDR values within each row. and the face colors indicate enrichment. Circles with black boundaries were statistically significant after FDR correction (α=0.05). The grey cells indicate missing results. Abbreviations: Ex – excitatory, In – inhibitory, OPC - oligodendrocyte progenitor cell. **B**, The union neuronal SCING gene regulatory network of three individual regulatory networks of the top KDs for the candidate genes with outgoing, incoming, and combined effects. Gene regulatory networks were constructed using SCING and neuronal scRNAseq data from different single cell Atlases including the Allen Brain Single Cell Atlas, the Mouse Cell Atlas (MCA), Tabula Muris, and Tabula Muris Senis. SCING networks were constructed from each dataset and the union network was used for key drier analysis. The larger, labeled circles indicate genes that were identified to be KDs, and smaller circles indicate candidate genes with the three effects. The direction of edges between genes indicates the regulatory relationship within SCING networks. For KDs, the color inside of each circle indicates from which candidate gene set the KD was identified, and multiple colors indicate the KD gene was a KD for multiple categories of candidate genes. The boundary color indicates whether the KD gene itself appeared as a candidate gene in any effect (or multiple effects). **C**, Enrichment analyses between the GWAS PD risk genes and our regulatory network genes. The outgoing, incoming, combined, and union represent the genes from different SCING networks (**Fig. S17-S20**, Fig. 7B). The *p* values were Bonferroni corrected with n=4.

### Key Driver analyses and GWAS enrichment analyses help to explore key driver genes

To further compare these candidate genes subsets, cell-type specific gene regulatory networks were constructed using SCING (Littman et al., 2023). Key driver (KD) analyses (Shu et al., 2016) were then performed to identify the network key drivers of the selective vulnerability related genes based on the network topology (see **Methods**), where key drivers are the network hub genes whose neighborhoods are over-represented by the vulnerability candidate genes. We firstly constructed the gene regulatory network for outgoing effect candidate genes and investigated their relationship based on the glial cell single cell RNA sequencing data (**Fig. S21, Supplementary Table - KD Analyses for outgoing effect (glial cell network)**), since outgoing effect candidate genes were enriched in glial cells (**Fig. 7A**). In order to deeper understand how these three candidate gene subsets work in the neuronal cell type, we built regulatory networks based on neuronal networks for outgoing (**Fig. S22**), incoming (**Fig. S23**), and combined effects (**Fig. S24**), respectively (see **Methods**) (see **Supplementary Table - KD Analyses (neuronal network))**. We merged these three individual networks (**Fig. S22-S24**) together to visualize the union gene regulatory network (**Fig. 7B**), which contained the top KD genes for all three effects.

Interestingly, KD genes within the same effect were closer to each other in the network (**Fig. 7B**), highlighting the differences between the three effects. The connections of outgoing and combined KD genes were denser than that of the incoming KD genes (**Fig. 7B**), demonstrating the more distant regulatory relationship of genes with incoming effect. To quantify this topological structure, we performed chi-square goodness of fit tests for KDs of each effect (**Fig. S25**). Our results showed that KD gene had closer relationship to genes from the same group than to genes from the other groups, and the KDs for the incoming genes had significantly fewer neighbor genes in the union network than the KDs for the outgoing and combined genes.

Finally, we analyzed the enrichment of candidate genes for PD risk genes from GWAS Catalog (Sollis et al., 2023), which showed that PD risk genes were significantly enriched in the outgoing regulatory network, the combined regulatory network, and the union of three regulatory networks (**Fig. 7C**, **Supplementary Table - GWAS_PD vs Regulatory Network Genes**). Overall, our gene regulatory network KD analyses further identified potential regulators and interactions among genes associated with selective vulnerability, pathological α-Syn spreading, and PD risk factors (**Fig. 7C**).

## Discussion

In this study, we provided a mathematical modeling approach and combined it with systems biology tools to discover potential risk genes for the spread of pathological α-Syn in PD. We investigated the nuanced directionality preference of pathological α-Syn spread, and successfully predicted its spreading along neuronal connectivity network in the central nervous system. Compared to previous studies (Dadgar-Kiani et al., 2022; Henderson et al., 2019a; Henderson et al., 2020; Mezias et al., 2020; Pandya et al., 2019), the current work considered the directionality preference of pathological α-Syn spreading as a parameter, allowing us to model directionally biased spread accurately. Our results suggest that the spread of pathological α-Syn in WT mice is predominantly retrograde but also has a small component of anterograde direction (**Fig. 2, Fig. S7A**). We also investigated the outgoing, incoming, and combined effects of gene expression on spreading directionality, which improved the model fitting performance over the global spread model and gene correlation models (**Fig. S11**) and again showed a similar predominantly retrograde direction as in the global spread model (**Fig. 5**). Our mathematical modeling therefore demonstrated that while α-Syn spread is predominantly retrograde, incorporating a minor anterograde component is important to improve the accuracy of data prediction in WT mice. More interestingly, we found that the spread directionality of pathological α-Syn was very different in *G2019S LRRK2* mice, which is the first report on how *G2019S LRRK2* affects the spread directionality preference. Overall, our current study has quantified the proportion of pathological α-Syn spreading for the first time, which gives a more nuanced picture of the directional preference of α-Syn than previously assumed (Dadgar-Kiani et al., 2022; Henderson et al., 2019a; Mezias et al., 2020; Pandya et al., 2019).

Although previous studies showed that the spread of pathological α-Syn followed the mesoscale connectome (Dadgar-Kiani et al., 2022; Mezias et al., 2020), it is not clear whether all connections contribute equally to pathological α-Syn spreading (Knox et al., 2018; Moon et al., 2014). We used bootstrapping method to demonstrate that pathological α-Syn propagated through the anatomic connections, and strikingly, we found that the spread of pathological α-Syn was primarily driven by the partial connectome composed of just the top 2% strongest connections (**Fig. 3**). We therefore propose the existence of the key subnetworks that drive the pathological spread for the first time (**Fig. 3, G-H**, **Fig. S8**), which allows us to localize the key brain regions and connections. This observation has also been validated by our model and the datasets from Henderson et al. (Henderson et al., 2019a), which showed that the minority of the connectome composed of the strongest connections could predict the pathological α-Syn spreading both in WT mice and *G2019S LRRK2* mice (**Fig. S9**). These findings enable us to assess the contribution of each of the strongest connections to pathological α-Syn spreading that can be explored with future experiments.

In addition to the connectome, we studied distinct effects of regional gene expression on pathological α-Syn spread, including the outgoing, incoming, and combined effects for the first time (**Fig. 4**; see **Methods**). Previously, only the outgoing effect has been modeled in mathematical models (Anand et al., 2022; Dadgar-Kiani et al., 2022; Henderson et al., 2019a). We investigated the case of four important PD risk genes, i.e., *Snca* (Henderson et al., 2019a), *Gba* (Henderson et al., 2020), and *Vps35* (Dadgar-Kiani et al., 2022), and *Gpnmb* (Diaz-Ortiz et al., 2022). Our study indicated that *Snca* and *Gpnmb* affected not only the presynaptic but also the postsynaptic regions, while *Gba* and *Vps35* mainly modulated the presynaptic sites (**Fig. 4, H-M, Fig. S15-S17**). Interestingly, none of these four genes are the top candidate genes that contribute to the selective vulnerability. These findings suggest that the genes modulating pathological α-Syn spreading may be distinct from those genes that are thought to be PD risk factors. In addition, it will be more cost-effective to use these theoretical models predicting the outgoing, incoming, or combined effects for individual gene before downstream functional studies, since each functional study may only focus on one partial step of the spreading. Overall, our study provides a new methodology for studying the trans-synaptic mechanisms of spread, especially the effects of the expression of specific genes on whether the source of the selective vulnerability emanates from the presynaptic or postsynaptic sites.

The Venn diagram (**Fig. S19**) showed a higher gene overlap ratio between genes with the outgoing effect and the combined effect than between pairs of gene sets with the other effects. Since the combined effect modeled the case that gene expression affected the presynaptic and postsynaptic regions equally, this result indicates that selective vulnerability is mainly driven by genes expressed at presynaptic rather than postsynaptic sites. It is also reflected in the fact that the outgoing and combined effects altered the Ave. CCC values much more than the incoming effect (**Fig. 4, K-L**). Given that the spread of pathological α-Syn is predominantly retrograde in WT mice (**Fig. 2**, **Fig. 5**), these results suggest that gene expressed at sites that receiving pathology rather than those that sending pathology predominantly modulates pathological α-Syn spread.

Our model allows us to rank the importance of genes by their model performance, showing a promising approach to explore PD risk genes from the perspective of pathological spreading (**Fig. 4, K-M**). For instance, we found that the top risk genes of combined effect (**Fig. 4M**, **Supplementary Table - Gene Results**) not only had great model performance but also were associated with the progression of pathological α-Syn, synucleinopathies, or the other neurodegenerative diseases (**Table 1**). Apart from these, genes from other effects may also be related to PD, such as the *Spp1* (149^th^ for incoming effect) (De Schepper et al., 2023) and *Dok5* (1^st^ for outgoing effect, and 2^nd^ for combined effect. **Table 1**).

**Table 1.** Relationship between top combined genes obtained from our model and evidences from previous studies.

| Gene name | Ranking | Related evidence | Reference |
| --- | --- | --- | --- |
| *Pde1a* | 1<sup>st</sup> | α-Syn toxicity | (Albert et al., 2022; Höllerhage et al., 2017) |
| *Dok5* | 2<sup>nd</sup> | Member of Dok gene family | (Miyoshi et al., 2017; Ueta et al., 2020) |
| *Mpped1a* | 3<sup>rd</sup> | Important in CA1 neurons | (Jiang et al., 2021; Zhong et al., 2020) |
| *Bmp3* | 4<sup>th</sup> | Member of Bmp gene family | (Vitic et al., 2021) |
| *Ctgf* | 5<sup>th</sup> | Dopaminergic neurodegeneration | (Chen et al., 2020; McClain et al., 2009) |
| *Slc2a3*,<br>*Slc2a13* | 6<sup>th</sup>-7<sup>th</sup> | Member of Slc2 gene family | (Aykaç and Sehirli, 2020; Vittori et al., 2014) |

The results of the GO analysis further summarized risk genes between pathological α-Syn spread and other related studies of synucleinopathies, and may explain the genetic co-morbidity mechanisms of PD and other neuropsychiatric disorders or neurological diseases. Besides learning, memory, and locomotory behavior (**Fig. 6B**, **Fig. 6D**), the candidate genes of our model were also related to biosynthetic and metabolic process of dopamine and catecholamine (**Fig. 6C**, **Fig. 6D**), which is in line with the known selective vulnerability of dopaminergic neurons in PD (Lundblad et al., 2012). Since the dopamine circuit is associated with the reward and decision-making system (Russo and Nestler, 2013), this result may reflect the co-morbidity between mood disorder and PD (Schrag and Taddei, 2017; Zhang et al., 2022). The genes were also related to ‘response to cocaine’ and ‘response to ischemia’ (**Fig. 6D**). The relationship between cocaine and PD are well documented (Fasano et al., 2008; Pregeljc et al., 2020; Villalba and Smith, 2013), particularly in the context of cocaine abuse (Curtin et al., 2015; Villalba and Smith, 2013) and the neurotoxicity (Bannon, 2005) leading to PD. Accordingly, since cocaine acts in dopaminergic neurons of ventral tegmental area and other brain regions (LaLumiere et al., 2012), they may share similar neural circuits as PD (Mash et al., 2003), and these neural circuits may be vulnerable to pathological α-Syn spread. Similarly, PD-related proteins can mediate secondary brain damage after cerebral ischemia (Kim and Vemuganti, 2017), and co-morbidity between stroke and PD has also been reported (Levine et al., 1992); our results therefore indicate a spreading-related mechanism underpinning the potential relationship between ischemia and PD as well. In summary, our results point to a promising bridge between PD, other neurological diseases, neuropsychiatric disorders, drug abuse, ion channels, and pathological α-Syn spread, and provide an insight into the risk genes involved in the mechanisms underlying their co-morbidity.

Cell type analyses further suggest that distinct cell-types play different roles in modulating pathological α-Syn spread, with microglial cells and neurons playing outsize roles at presynaptic sites and postsynaptic sites, respectively (**Fig. 7B**). However, there was no significant cell-type preference in the combined effect (**Fig. 7B**); given that the combined effect blurs the contributions of outgoing and incoming effects, this result emphasizes that studying the impact on outgoing and incoming connections separately is required to disentangle the cell types involved in mediating each process.

To assess the interactions among the candidate genes involved in α-Syn spreading, we modeled gene regulatory networks and our key driver analyses revealed obtained KD genes which are potential regulators of the candidate genes. These KD genes may play important roles in regulating PD or synucleinopathies. Among the KD genes whose network neighborhoods showed top fold enrichment rankings for candidate genes with multiple effects are *Mpped1* and *Ctgf* discussed above, as well as *Dlgap2*, whose network neighborhood has highest fold enrichment for genes with outgoing and combined effects and is associated with age-related cognitive decline (**Fig. S22-S24**, **Supplementary Table - KD Analyses (neuronal network)**) (Ouellette et al., 2020). By identifying the relationships between KD and candidate genes, KD analyses also allow us to elucidate the regulatory networks involved in mediating the spreading of α-Syn pathology. As expected, the GWAS enrichment analyses showed that the combined and union regulatory networks had higher enrichment for PD risk genes (**Fig. 7C**), demonstrating that the mechanisms by which these genes contribute to the risk of developing PD include the mediation of network spreading of α-Syn.

Another significant outcome of the current study is that we present a pipeline for investigating the mechanisms of pathological progression from mathematical modeling and systems biology perspectives (**Fig. 6**, **Fig. 7**). Future work includes applying our mathematical models to PD patient imaging data, which may become a promising tool for diagnosis and disease treatment at early stage. Although there have been promising recent developments towards direct, in vivo imaging of α-Syn using positron emission tomography (PET) (Xiang et al., 2023), no publicly available datasets are currently available for analyzing the spatiotemporal deposition of α-Syn in human subjects. Nevertheless, previous mathematical modeling by our group and others has been employed to explain the progression of neurodegeneration as assessed by deformation-based morphometry (DBM) in PD patients (Freeze et al., 2019; Pandya et al., 2019), and also explained the patterns of neurodegeneration of AD and other tauopathies (Iturria-Medina et al., 2014; Raj et al., 2015; Weickenmeier et al., 2018), as well as amyotrophic lateral sclerosis (ALS) (Pandya et al., 2022). Recently, synaptic oligomeric tau was directly observed in the brains of AD patients (Colom-Cadena et al., 2023), providing further motivation for using mathematical models in prion-like diseases. Combined with the regional gene expression data in humans provided by the AIBS (Hawrylycz et al., 2012), our model will provide a critical tool to analyze the mechanisms of gene-mediated pathological protein spread throughout the brain in other neurodegenerative diseases.

We note that the current study has several limitations, which we plan to further investigate experimentally in the future. While the AGEA (Lein et al., 2007) remains the most expansive resource for spatial gene expression data in the mouse brain, we recognize that gene expression levels may be altered with the disease progression after PFF injection, which the AGEA does not capture. Consequently, we can only evaluate the effect of *baseline* gene expression on pathology spread, as many previous studies have done (Anand et al., 2022; Cornblath et al., 2021; Dadgar-Kiani et al., 2022; Henderson et al., 2019a; Henderson et al., 2020). We plan to perform future functional analyses to experimentally validate the genes identified by our model. Additionally, our study was limited by the current resolution of Allen Mouse Brain Atlas (AMBA) and the mesoscale connectome data (Knox et al., 2018; Moon et al., 2014) consisting of 410 brain regions; this may have led us to overlook pathology spread between potentially important subregions as well as local neural networks. We therefore plan to perform in vitro experiments to investigate spread processes at finer spatial scales. Besides, our model does not incorporate the effects of degeneration, since we only studied the progression at early stages of the disease, and previous studies have demonstrated there are few brain regions that show significant cellular degeneration before 6 MPI (Henderson et al., 2019a; Rey et al., 2018). However, degeneration may play a more significant role at later stages of disease, and therefore we will conduct more experiments to assess α-Syn pathology beyond 6 MPI and update our model accordingly.

## Supporting information

Supplementary file (ZIP): Includes all Supplementary Tables and Files.

## Acknowledgements

This work was supported by NIH/NINDS R01-NS128964, NIH/National Center for Advancing Translational Science (NCATS) UCLA CTSI Grant Number UL1TR001881, and the Noble Family Innovation Fund (to C.P.); NIH/NIA RF1AG062196 and AG072753 (to A.R.); NIH/NINDS R01-NS111378 and NS117148 (to X.Y.). The authors also thank Zhaoxu Liu ‘Slandarer’ for developing the open-source color palette toolbox and correlation bubble chart visualization toolbox for MATLAB (https://www.mathworks.com/matlabcentral/profile/authors/18192500).

## Author Contributions

Y.L. conceived and designed the experiments, collected data, performed modeling and coding, analyzed and visualized the results, wrote the original draft, and edited the draft. J.T. conceived and designed the experiments, performed modeling and coding, analyzed and visualized the results, wrote and edited the manuscript. J.D. performed gene network and cell-type enrichment analyses, and visualized the results. N.W. performed GWAS-PD enrichment analyses and visualized the results. C.L. and S.K. performed annotations and quantifications for brain sections, collected data, analyzed the results. C.A. collected data. J.T. and M.C. curated multiple single cell Atlases and constructed cell-type specific SCING networks. C.L. and B.L. analyzed and visualized the results. Y. S. revised the manuscript. X.Y. supervised, conceived and designed the systems biology analyses, and edited the manuscript. A.R. supervised the study, conceived and designed the experiments, wrote and edited the draft. C.P. planed and supervised the study, conceived and designed the experiments, interpreted the data, wrote and edited the draft. All authors reviewed and approved the manuscript.

## Data and Code Availability

All source code and data files are available on GitHub (https://github.com/Yuanxi-Li/PFF_Progression_Model) upon reasonable request.

## Methods

### Animals

Two-to-three-month-old C57BL6/C3H wild-type mice were purchased from the Jackson Laboratories (Bar Harbour) for the stereotaxic injection experiments. All breeding, housing, and experimental procedures were performed according to the NIH Guide for the Care and Use of Experimental Animals and approved by the University of Pennsylvania Institutional Animal Care and Use Committee (IACUC).

### *In vitro* α-Syn preformed fibrils generation

Full-length mouse α-Syn (1-140) monomers were expressed in BL21 (DE3) RIL cells and purified referring to the previous protocols (Giasson et al., 2001). Mouse preformed fibrils (PFFs) was generated by diluting the above α-Syn monomers to 5mg/ml in sterile Dulbecco’s PBS (Cellgro, Mediatech; pH adjusted to 7.0), and then incubating it at 37°C with constant agiration at 1,000 r.p.m. for 7 days. The sedimentation test and thioflavin T-binding assay were used to verify α-Syn fibrillization (Volpicelli-Daley et al., 2014).

### Stereotaxic injection of mouse α-Syn PFFs

Two-to-three-month-old C57BL6/C3H wild-type mice were anaesthetized with ketamine hydrochloride (100 mg/kg), xylazine (10 mg/kg) and acepromazine (0.1 mg/kg). The amount of 6.25μg mouse PFF in 2.5μl Dulbecco’s PBS was stereotaxically injected into the dorsal striatum (caudoputamen; coordinates: 0.2 mm relative to bregma, +2.0 mm from midline, +3.2 mm beneath the surface of skull) with 10-μL syringes (Hamilton) at a rate of 0.4 μL per min. Mice were euthanized at 3 and 6 months-post-injection.

### Immunohistochemistry

Mice were transcardially perfused with PBS. The brain and spinal cord were removed and fixed in 70% ethanol (in 150mM NaCl, pH 7.4) overnight. After fixation, brains were embedded in paraffin blocks and the immunohistochemistry was performed as previously described (Duda et al., 2000; Luk et al., 2012b). Phosphorylated at Ser 129 (pS129, 81A, 1:10000 dilution) antibodies were used for misfolded α-Syn proteins. Staining sections were scanned by a Perkin Elmer Lamina scanner at 20× magnification. Sample size: n=4 for the three-months-post-injection mice, n=3 for the six-months-post-injection mice.

### Quantitative pathology

All section selection, annotation, and quantification were performed by mouse neuroanatomists who were blinded to the treatment. For the quantification of the α-Syn pathology, coronal slices were selected to closely match the following nine bregma levels: +4.28 mm, +2.10 mm, +0.98 mm, −0.22 mm, −1.22 mm, −2.18 mm, −2.92 mm, −3.52 mm, and −4.48 mm. For each bregma level of an individual mouse, 2 slices were selected as the technical replicates. The digitized images were uploaded into the QuPath software to allow annotation and quantification of the percentage area occupied by α-Syn pathology. Standardized annotations were drawn to allow independent quantification of 196 grey matter regions throughout the brain. Each set of annotations was hand-drawn onto the desired sections to match the designated brain regions. After annotations of all brains, analysis algorithms were applied to all stained sections, and data analysis measures for each region were recorded.

One analysis algorithm was applied to the tissue, but it varied depending on the level of background staining observed in the mice. The algorithm detected a total signal above a minimum threshold, which was determined to most likely not include any background signal. Specifically, the analysis included all signal that was above a 0.5 or 0.7 optical density threshold, based on the analysis algorithm used. Additionally, the analysis tool was developed with a resolution of 0.49 microns per pixel and a smoothing sigma of 0.5. The signal was then normalized to total tissue area.

Finally, in order to match our quantitative data to the connectome data for subsequent modeling work, we merged and added some brain regions. We eventually obtained the pathological dataset containing 410 grey matter regions, with 205 regions bi-hemispheric symmetry. Among these, data of 156 brain regions were derived from the quantification of pathological immunohistochemistry. The value of each brain region represents the regional pathological burden (the area of pathology over the area of the whole region). Since the amounts of pathology between different regions could vary by a factor of 1∼1,000, we took the logarithm transformation for visualizing the quantitative data at 3 and 6 MPI averaged within its group.

### Mathematical models

#### Global spread model

The α-Syn pathological progression at early stage was considered as the sum of the following two effects (Anand et al., 2022; Peng et al., 2018; Peng et al., 2020; Rey et al., 2018): 1. Pathology spread. If two brain regions were structurally connected, the pathology could be spread from one region to the other. The spread can be driven by the concentration gradient (diffusion) or against the gradient (active transport). From the cellular level, the spread directionality can be fully anterograde (only from presynaptic membrane to postsynaptic membrane), fully retrograde (only from postsynaptic membrane to presynaptic membrane) or directionally biased. 2. Amplification and clearance. The pathological α-Syn seeds will induce α-Syn monomers to generate new misfolded α-Syn aggregates via template amplification, and the misfolded α-Syn aggregates can be removed by autophagy and lysosomal degradation. The total amount of the pathology in the whole network were determined by the overall effects of the amplification and clearance. We did not consider the degeneration process, because although some papers have reported the degeneration of α-Syn pathologies after 8 MPI (Dadgar-Kiani et al., 2022), pronounced degeneration before 6 MPI in the whole brain has not been observed (Henderson et al., 2019a; Rey et al., 2018).

In our first set of analyses, we augmented the Nex*is:*global model (Anand et al., 2022), and applied it to model the two process in the brain networks. The original Nexis model has shown a great performance on predicting the spread of pathological Tau (Anand et al., 2022). Compared to the previous Network Diffusion Model (NMD) (Henderson et al., 2019a; Raj et al., 2012), the Smoluchowski network model (SNM) (Bertsch et al., 2021; Fornari et al., 2020; Raj et al., 2021), the original Nex*is*:global (Anand et al., 2022), our augmented model was able to simultaneously capture the seed scaling, the global amplification or clearance effect, the diffusion effect, and the directionally biased spread of pathology, whereas the others were only modelling parts of the above factors, especially ignoring the possibility of directionally biased spread.

We first modeled the spread component. Let *X* ={*x_i_* | *i* = 1, 2,…, 410} be the area fractions of the α-Syn pathology in our 410 grey matter regions, and *C* = {*c_ij_* | *i*, *j* = 1, 2,…, 410} be the weighted adjacency matrix that defines the axonal projection strength from brain region *i* to *j* (connectome adjacency matrix) (Knox et al., 2018; Moon et al., 2014). First, we consider the case where spread was directional and fully anterograde. For each individual brain region *i*, pathology spreads from region *j* to *i* along projection *c _ji_*, which increases the pathology burdens of region *i* ; meanwhile, pathology also spreads from region *i* to *j* along projection *c_ij_*, which decreases the pathology burdens of region *i*. Therefore, the amount of pathologies changed by spread can be described as the following ordinary differential equation (ODE):

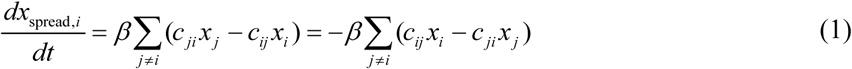

where *β* was the global diffusivity rate to modulate the velocity of the pathology diffusion, *t* ∈ *N* denoted the month post injection (MPI).

Then, we defined the amplification and clearance process using a global parameter *α*. *α* was the overall rate of the amplification and clearance effects. We let the positive *α* denote the overall amplification effect. For each brain region *i*, the rate of change of pathology through amplification and clearance was given by:

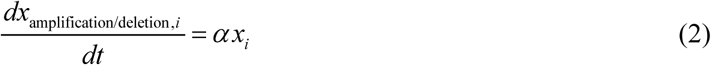

The fully anterograde global spread model, then, was governed by the sum of these two equations:

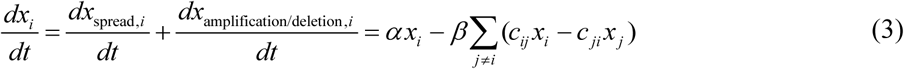

We defined the diagonal operation 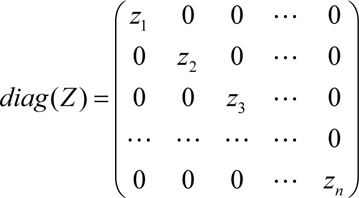, and let *columnsum* denotes the sums of every column of a given matrix. Then, equation (3) had an analytical solution: *X̂* (*t*) = *e*(*diag* (*α*)−*β L*)*t X*. Here, *X̂* (*t*) was the vector of the predicted pathology, *L* was the Laplacian operator of the connectome adjacency matrix, where *L* = *L*_anterograde_ = *diag*(*columnsum*(*C*′)) − *C*′ and 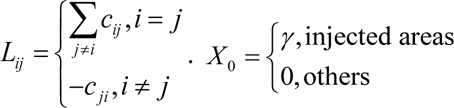 was the seed scaling vector that described the amount of the injected pathological seeds at 0 MPI, which were 0 for not-injected regions and *γ* ∈ (0,1] injected regions.

Similarly, for the fully retrograde diffusion, there was only one different term:

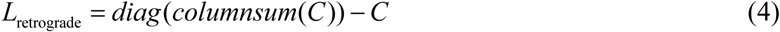

In order to accommodate unequal bidirectional spread, we introduced the parameter *s*, which was bounded between 0 and 1. By convention, a value of 0 indicated fully anterograde spread, a value of 1 indicated fully retrograde spread, and a value of 0.5 indicated unbiased spreading. We therefore generalized the spread equation (1) as follows:

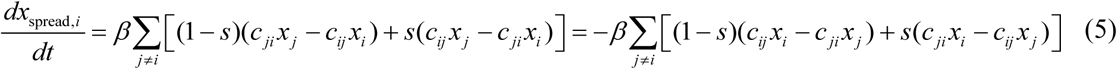

The general Laplacian operator, *L̂*, was therefore defined as:

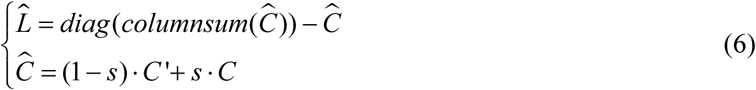

As before, this equation had the following analytical solution:

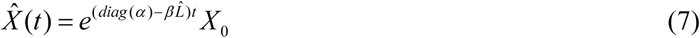

#### Spread model with the effect of gene expression

In addition to the connectomic spread and the global amplification and clearance effects, it has been reported that the cell type densities (Peng et al., 2018) and regional gene expression (Dadgar-Kiani et al., 2022; Henderson et al., 2019a; Henderson et al., 2020) also play very important roles in the pathology progression; this is known as the selective vulnerability of different brain regions imparted by genes. To investigate this effect, and especially to identify the candidate genes that were associated with the pathological spread, we explored the effects of the 3,855 individual genes from the coronal series of the AGEA (Anand et al., 2022; Lein et al., 2007). We modified the Nex*is*:microglia model (Anand et al., 2022) explored previously to accommodate the differential effects of outgoing and incoming connections as follows.

Let 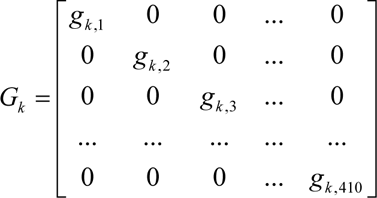, where *g_k_*_,*i*_ was the gene expression level of gene *k* on the brain region *i*. Then we define *gene-mediated connectivity matrix*, *C* ⊗ *G_k_*, which can take different functional forms depending on whether the effect on presynaptic or postsynaptic sites is being considered. Using this nomenclature, previous studies have defined *C* ⊗ *G_k_* = *G_k_* · *C*, and therefore have only explored the gene mediation of pathology spread along outgoing connections (**Fig. 4B**) (Anand et al., 2022; Dadgar-Kiani et al., 2022; Henderson et al., 2019a). Biologically, these models only investigate the selective vulnerability from the gene expression of presynaptic regions (**Fig. 4C**). In order to generalize the model to also accommodate modulation from incoming regions alone or equally from both the outgoing and incoming connections simultaneously, we have considered the following three forms of *C* ⊗ *G_k_* (**Fig. 4, A-D**):

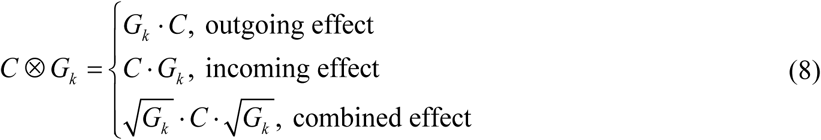

Therefore, the direction-biased spreading of the gene-connectome matrices were updated as *Ĉ_Gene_* = (1 − *s*) · (*C* ⊗ *G_k_*)′ + *s* · (*C* ⊗ *G_k_*), and its related Laplacian matrices were then updated as *L̂_Gene_* = *diag*(*columnsum*(*Ĉ_Gene_*)) − *Ĉ_Gene_*. Then, the solution can be calculated by

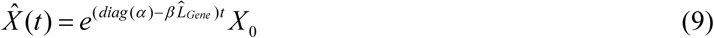

#### Model fitting with the quantitative pathological data

We took the average of the 3 and 6 MPI’s quantitative pathological data respectively, and fitted the above models over *t* ∈{3,6}. We picked the seed scaling parameter *γ*, the overall effect of global amplification or clearance *α*, the global diffusivity rate *β*, and the directionality parameter *s*, which maximized the model fitting, defined as the best concordance correlation coefficients (CCC) (Lawrence and Lin, 1989) between the quantitative pathology vector log_10_ ( *X* (*t*)) and the predicted pathology vector log_10 (_ *X̂* (*t*)_)_ for all non-zero values of *X* (*t*), averaged over *t* ∈{3,6}. Compared to Pearson’s correlation coefficient method as a loss function, CCC can capture the scale characteristics (Lawrence and Lin, 1989) and be used to fit the seed scaling parameter *γ*. We took the logarithm transformation, since the scale of the quantitative pathology data in different regions could differ by a factor of 1-1,000. According to the above models, the parameters had different bounds limitations. Using the MATLAB (R2022a, MathWorks) function ‘fmincon’, we fit the parameters of the model using the following cost function:

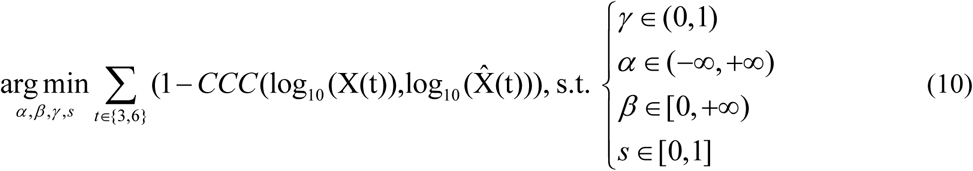

The model ran successfully for all genes in the outgoing or incoming effect models, but 60 genes failed to be fitted in the combined effect due to the error with matrix computations (3795 out of 3855 genes retained, **Supplementary Table - Gene Results**).

### Robustness testing of mathematical models

Bootstrapping was used to assess robustness. To prove that pathology progression was driven by the mesoscale connectome but not by randomly generated matrices, we tested the global spread model with 1,000 null connectomes whose elements were sampled from the uniform random distribution *U*_[0,1]_. We also explored the effect of randomly permuting the actual connectome weights, creating a different set of 1,000 null connectomes. To investigate whether the partial connectome was able to explain the pathology progression, and also to explore the characteristics of the key partial connectome, we fit the global spread model using connectomes with specified percentages of connections removed at random, with each 5% increment sampled 1,000 times.

We also tested the robustness of the genes using the 500^th^ genes of each spread effect (*Large* for the outgoing effect, *Socs6* for the incoming effect, and *Tbc1d14* for the combined effect) as test cases. The bootstrapping was repeated 1,000 times by randomly permuting the elements of the gene expression vectors.

The random seeds (‘rng’ function in MATLAB software) were incorporated to ensure that the results of bootstrapping were reproducible.

### Model validation using other datasets

In order to validate our findings of spreading directionality and key connectomes, we ran our model using datasets from Henderson et al. (Henderson et al., 2019a). Similar to our experiment, the amount of 5μg mouse α-Syn PFFs in 2.5μl Dulbecco’s PBS were injected into the dorsal striatum of the C57BL/6J non-transgenic mice (NTG mice, same as the wild-type mice in our study) and the B6.Cg-Tg(Lrrk2*G2019S)2Yue/J *G2019S LRRK2* transgenic mice (*G2019* mice). Mice were euthanized at 1, 3 and 6 months-post-injection (NTG mice: n=4 for 1 MPI, n=6 for 3 MPI, and n=6 for 6 MPI. *G2019* mice: n=6 for 1 MPI, n=6 for 3 MPI, and n=7 for 6 MPI), and all mice were initially stained with Syn506 for quantification of pathology. After brain annotation and quantification, α-Syn pathology distribution data and a connectome of 116 annotated brain regions (13,456 connections in total, denoted as the ‘masked’ connectome) were used to model the pathological progression according to the above global spread model. Note that compared with our whole connectome (410 brain regions with 168,100 connections in total), the ‘masked’ connectome in Henderson et al. (Henderson et al., 2019a) only represents a subset of all connections in the brain. The detailed dataset and experimental design can be found in Henderson et al. (Henderson et al., 2019a) and on GitHub (https://github.com/ejcorn/connectome_diffusion).

### Systems biology analyses

#### Candidate genes selection

After fitting the above models, we ranked the importance of the genes for their outgoing, incoming, and combined effects, separately. Any individual gene related to a specific spread effect will have an Ave. CCC values for its best fitting, and the genes which had better Ave. CCC values than that of the global spread model were considered as the potential genes which affect the spread positively. We picked the top 500 genes from the potential gene lists for each effect group as the candidate genes. We then performed gene ontology analyses, cell type analyses, and gene regulatory network key driver analyses.

#### Gene ontology analyses

Gene ontology (GO) analyses were performed in PANTHER to study the enrichments in biological process, molecular function, and cellular component of the candidate genes (PANTHER Overrepresentation Test (Released 20221013); GO Ontology database DOI: 10.5281/zenodo.7709866 Released 2023-03-06; http://geneontology.org/) (Mi et al., 2019). For each GO catogory of any specific spread effect, we visualized the results sorted by fold enrichment that were statistically significant after Bonferroni correction.

#### Cell-type analyses

Marker genes for neuronal cell types (excitatory and inhibitory) and glial cell types (astrocyte, microglia, oligodendrocyte, oligodendrocyte progenitor cell, and ependyma) were used to test for enrichment in the candidate genes of the incoming, outgoing, and combined effects. Excitatory and inhibitory neuronal marker genes were collected from the study on cell type compositions across the entire mouse brain (Langlieb et al., 2023). Distinguishing markers from different excitatory (2426 cell types) and inhibitory (2242 cell types) neurons were combined into excitatory and inhibitory marker gene lists. As glial cell marker genes are more similar across brain regions, an internal single-cell RNA-sequencing mouse dataset (See **Supplementary File - Brain_cell_type_markers** for details) was used to retrieve marker genes for astrocyte, microglia, oligodendrocyte, oligodendrocyte precursor cell, and ependyma consistent between the hippocampus and hypothalamus (average log2FC > 2 compared to other cell types for each tissue) which captured canonical glial cell type marker genes. Marker gene lists were subset to the genes used for testing of the model. Hypergeometric test was used to retrieve the significance of enrichment, and *p*-values were Bonferroni adjusted. The background population size was the number of genes (3855) tested in the model.

#### Key driver (KD) analyses and regulatory network

We used SCING (Littman et al., 2023) to build gene regulatory networks for candidate gene subsets. We used the default hyperparameters in SCING that included all genes, 10 principal components, 0.7 subsampling proportion, 100 neighbors per gene, and 500 target supercells. Single cell transcriptomics data from Tabula Muris (Schaum et al., 2018), Tabula Muris Senis (2020), and Mouse Cell Atlas (Han et al., 2018), with networks built on the Allen 10X dataset (Yao et al., 2021) from the hippocampus, visual cortex, somatosensory cortex, and primary motor cortex brain regions was used for outgoing effect candidate genes, since outgoing effect candidate gene was enriched in glial cells. Then, to study the gene regulatory relationship in neuronal network, one neuronal SCING network was built based on the neuronal single cell transcriptomics data from Allen Brain Atlas (Daigle et al., 2018), Tabula Muris (Schaum et al., 2018), Tabula Muris Senis (2020), and Mouse Cell Atlas (Han et al., 2018), for each candidate gene of three effects (outgoing, incoming, combined) to the neuronal cell types and the resulting networks were combined together as a union network with the data source information retained. This union network along with the candidate genes of the incoming, outgoing, and combined effects were used as inputs into key driver analysis (KDA) (Shu et al., 2016) using a search depth of 1 and not considering direction to identify KDs whose neighborhood subnetworks were enriched for the candidate genes of each category (incoming, outgoing, combined) based on a Chi-like statistics. We called key drivers with an FDR threshold of less than 0.05.

To test whether KDs for the candidate genes of the same effect category (incoming, outgoing, or combined) are more connected than KDs for candidate genes of different categories, we performed chi-square goodness of fit tests for each KD effect, with the observations being the number of the directed edges from a source KD node of each effect to its respective target KD nodes. As each node could fall under multiple effects, we used two effects for each edge: target was of the same effect or not. The expected number of edges for each KD to the same source type was calculated by

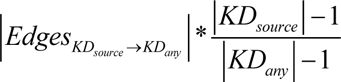 based on the number of edges for each effect category found in the networks and the expected probabilities for each KD based on the gene metadata if edges selection was independent. The remaining edges were expected to not be of the source KD type.

#### GWAS-PD enrichment analyses

We performed enrichment analysis on the genes in each network (incoming, outgoing, combined, and the union network of all three) for over-representation of PD candidate risk genes from the GWAS Catalog (Sollis et al., 2023) using hypergeometric tests (Paczkowska et al., 2020) with Bonferroni correction for n=4.

### Software and statistics

All statistical tests were two-tailed, and the significance level was set to *α* = 0.05 unless otherwise noted. Normality was assessed using the Shapiro-Wilk test. Student’s t-test (for normally distributed data) or Wilcoxon rank-sum test (for non-normally distributed data) was used for comparing pathology between 3 and 6 MPI for each quantified brain region. Bonferroni correction and the false discovery rate (FDR) were used for multiple comparisons in systems biology analyses. Chi-square test was used for comparing the actual number of the edges with the expected number after KD analyses. All of the *p* values were adjusted accordingly. All statistical analyses and visualization were performed mainly with MATLAB, Graphpad Prism (8.0.2, GraphPad Software), R (4.1.3), and Python.

## Supplementary Figures

**Figure S1.**
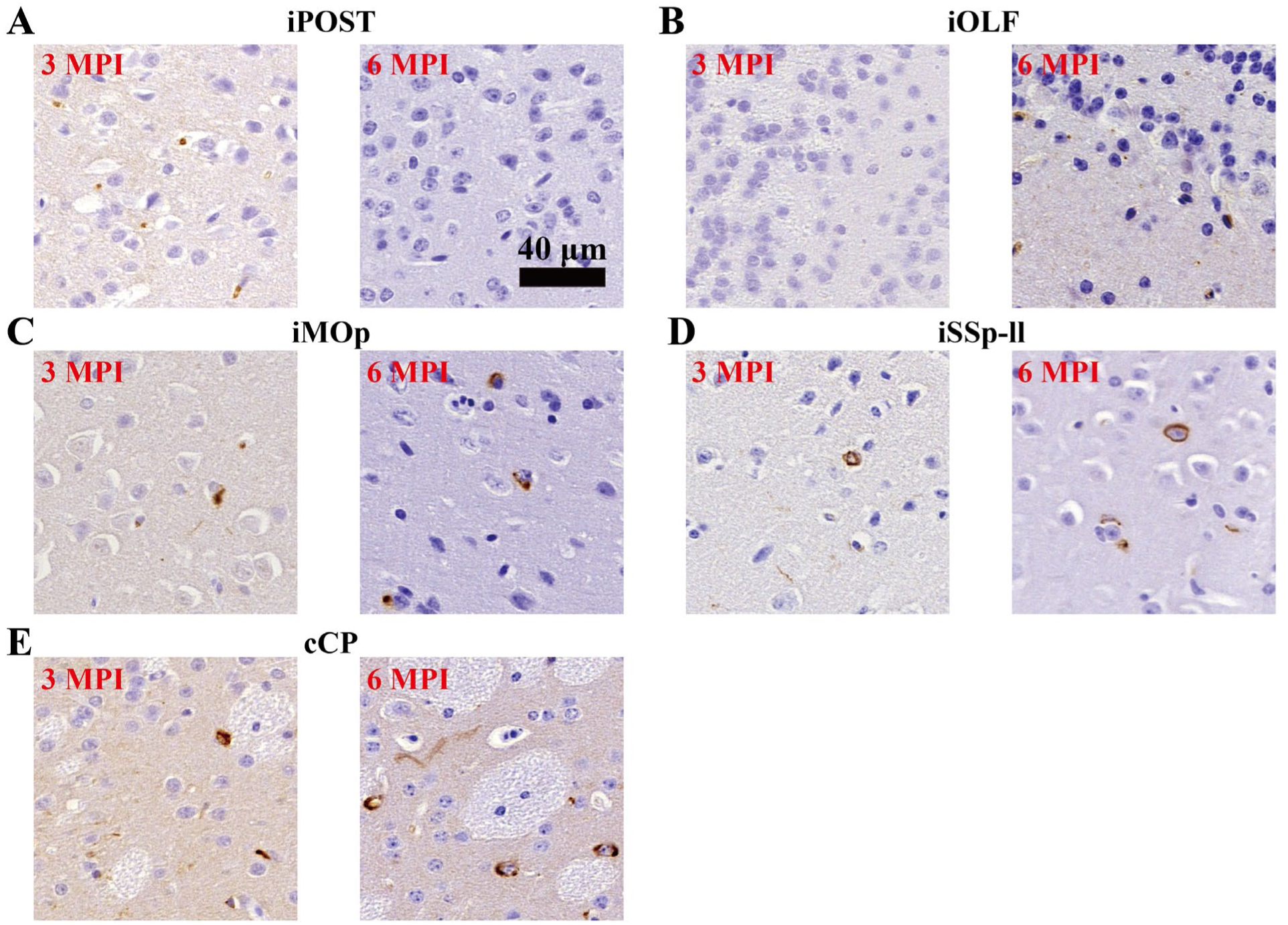
Representative quantitative pS129 α-synuclein pathology images of selected regions at 3 and 6 MPI. **A**, iPOST region, showing a decrease trend from 3 to 6 MPI. **B-E**, Regions with statistically significance between 3 and 6 MPI after FDR correction. **B**, iOLF region. **C**, iMOp region. **D**, iSSp-ll region. **E**, cCP region. All scales were the same and the scale bar was shown in **A**. Scale bar: 40 μm.

**Figure S2.**
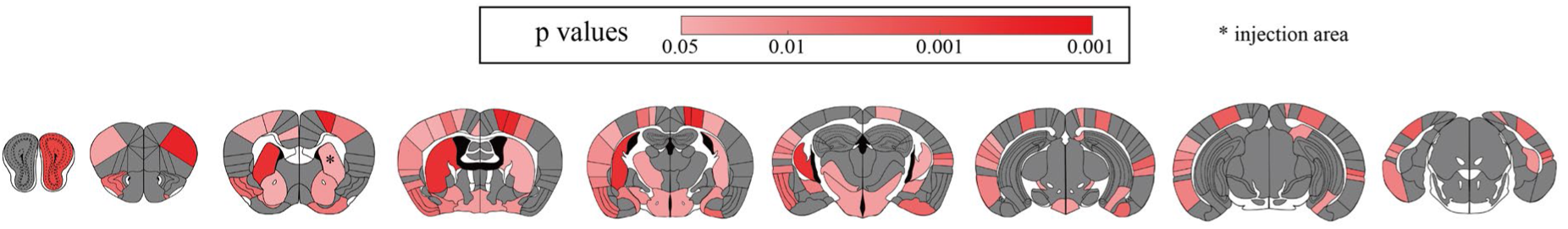
Heatmap showing regional *p* values with statistically significant (*p*<0.05) differences between 3 and 6 MPI. Warmer colors represented the differences between 3 and 6 MPI were more statistically significant (*, injection area).

**Figure S3.**
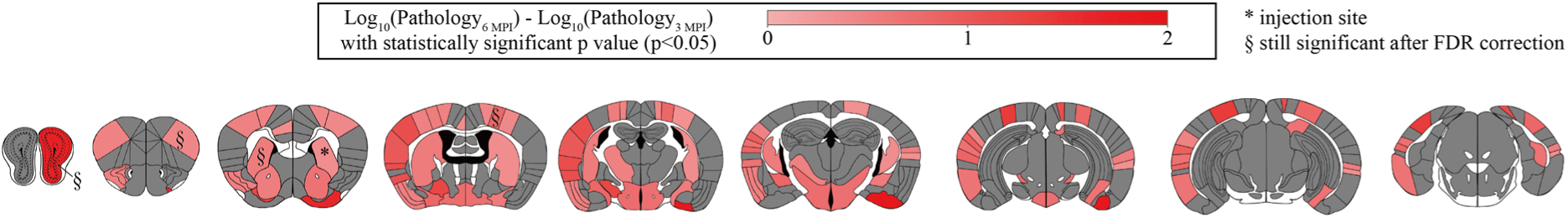
Differences of regional pathology burdens between 3 and 6 MPI, only displaying the results of statistically significant regions. All the statistically significant regions showed upward trends of pathology. iOLF, iMOp, iSSP-ll, and cCP were still significant after false discovery rate (FDR) correction. *, injection area. §, regions that were still significant after FDR correction.

**Figure S4.**
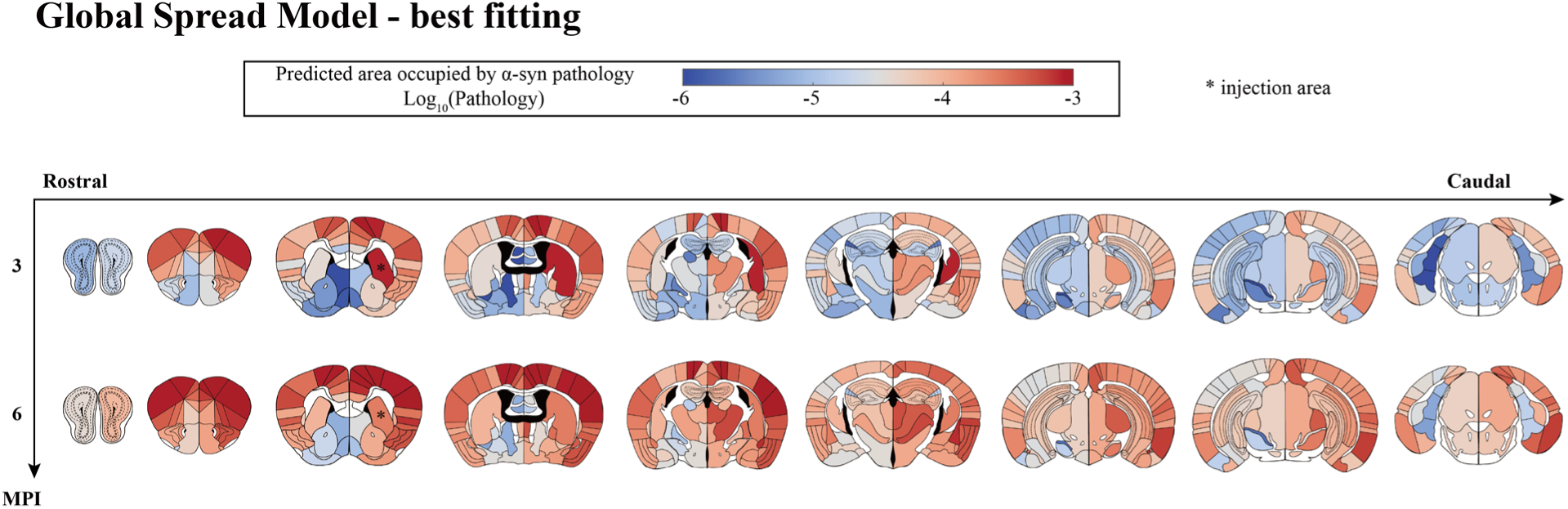
Heatmap of the regions affected by α-Syn pathologies calculated by the best model fitting of global spread model (log10-transformed; *, injection area).

**Figure S5.**
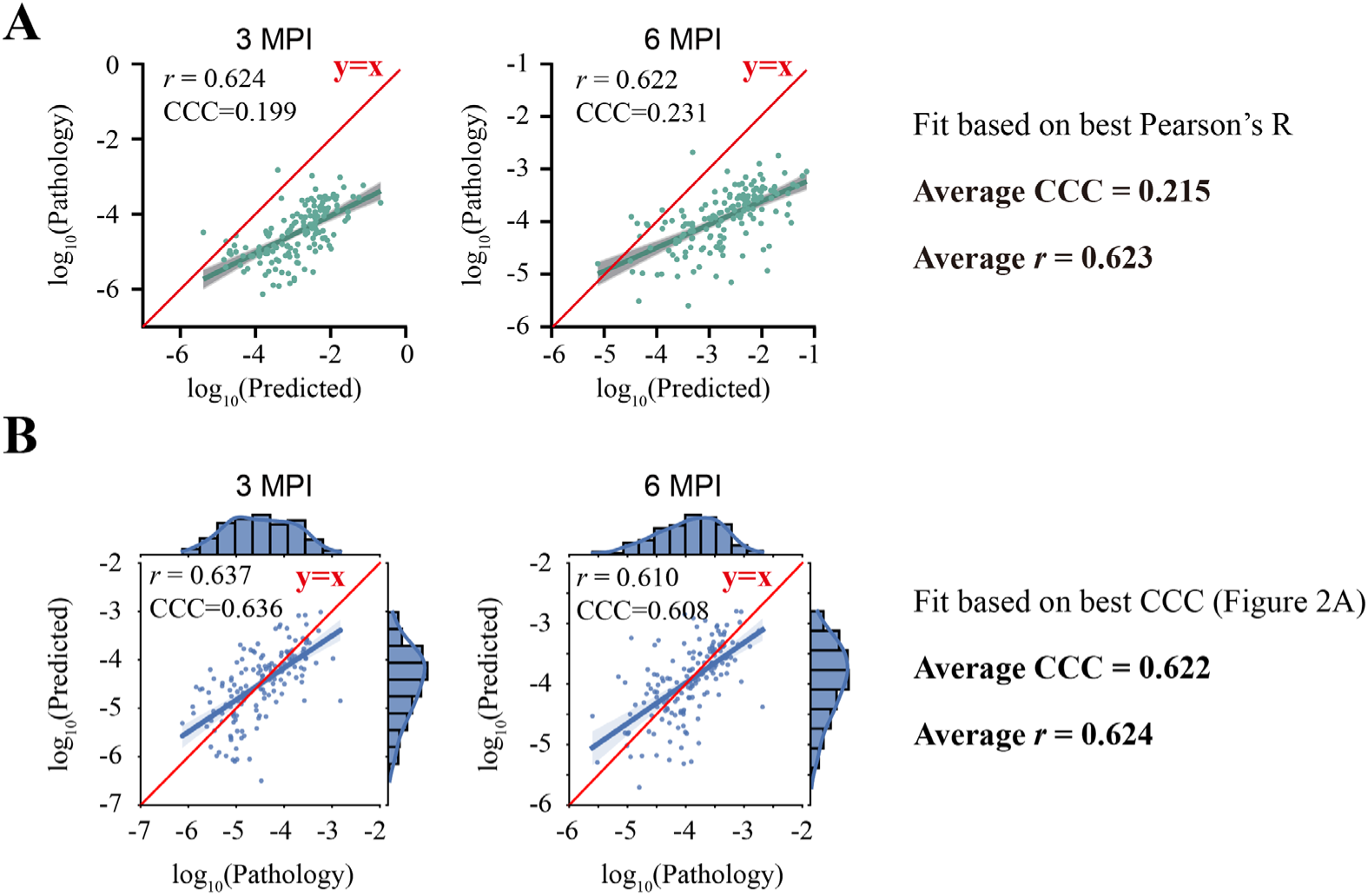
Comparison between the model fitting using Pearson’s R (**A**) and CCC (**B**, which reproduces Fig. 2A). The closer the best-fit line is to y=x (red line), the better the model captures scale of the observed data.

**Figure S6.**
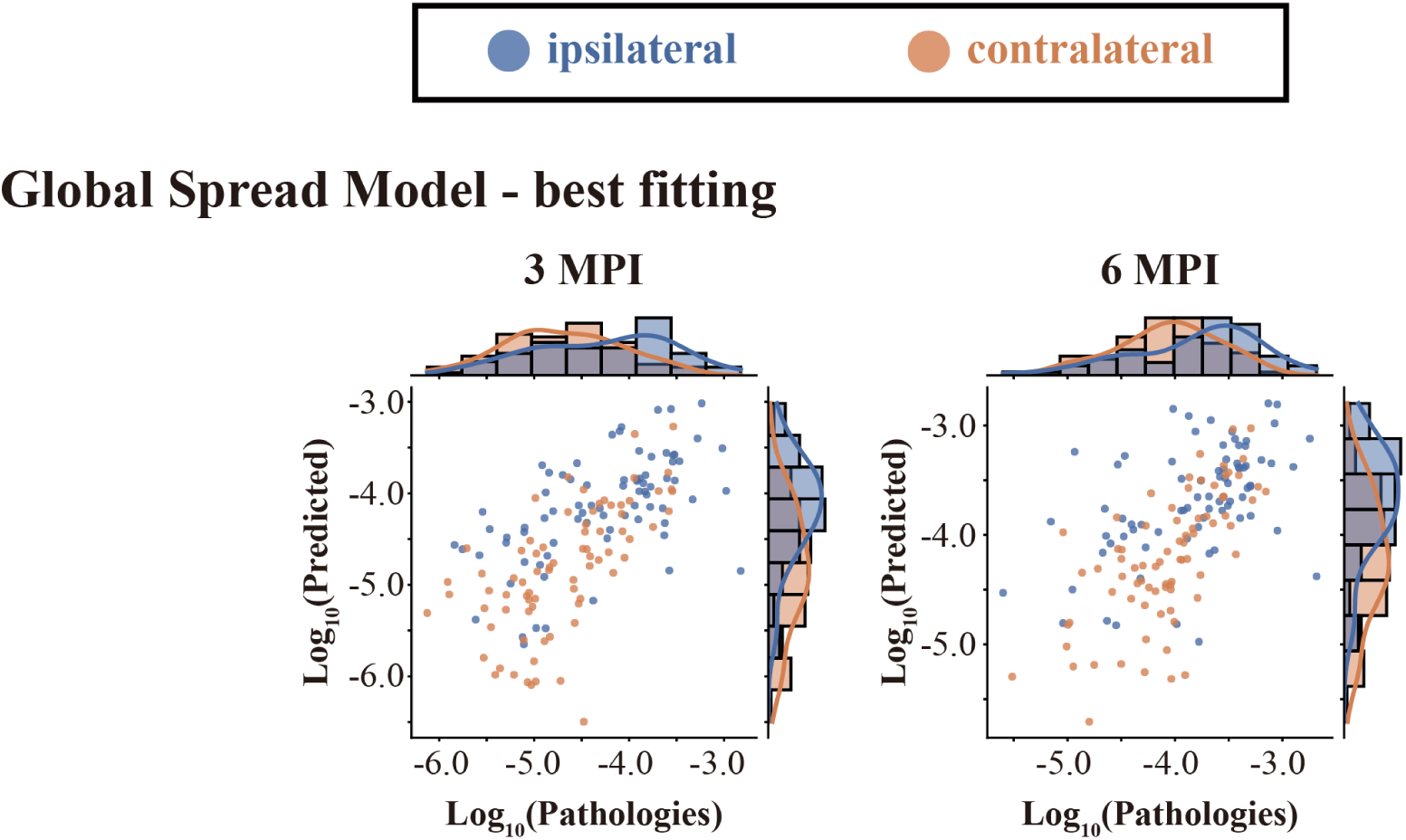
The global spread model accurately captured the feature that the ipsilateral hemisphere had more pathology than the contralateral hemisphere both at 3 and 6 MPI. Regions ipsilateral to the injection site are shown in blue, while contralateral regions are shown in orange, with histograms of the univariate distributions shown along the axes.

**Figure S7.**
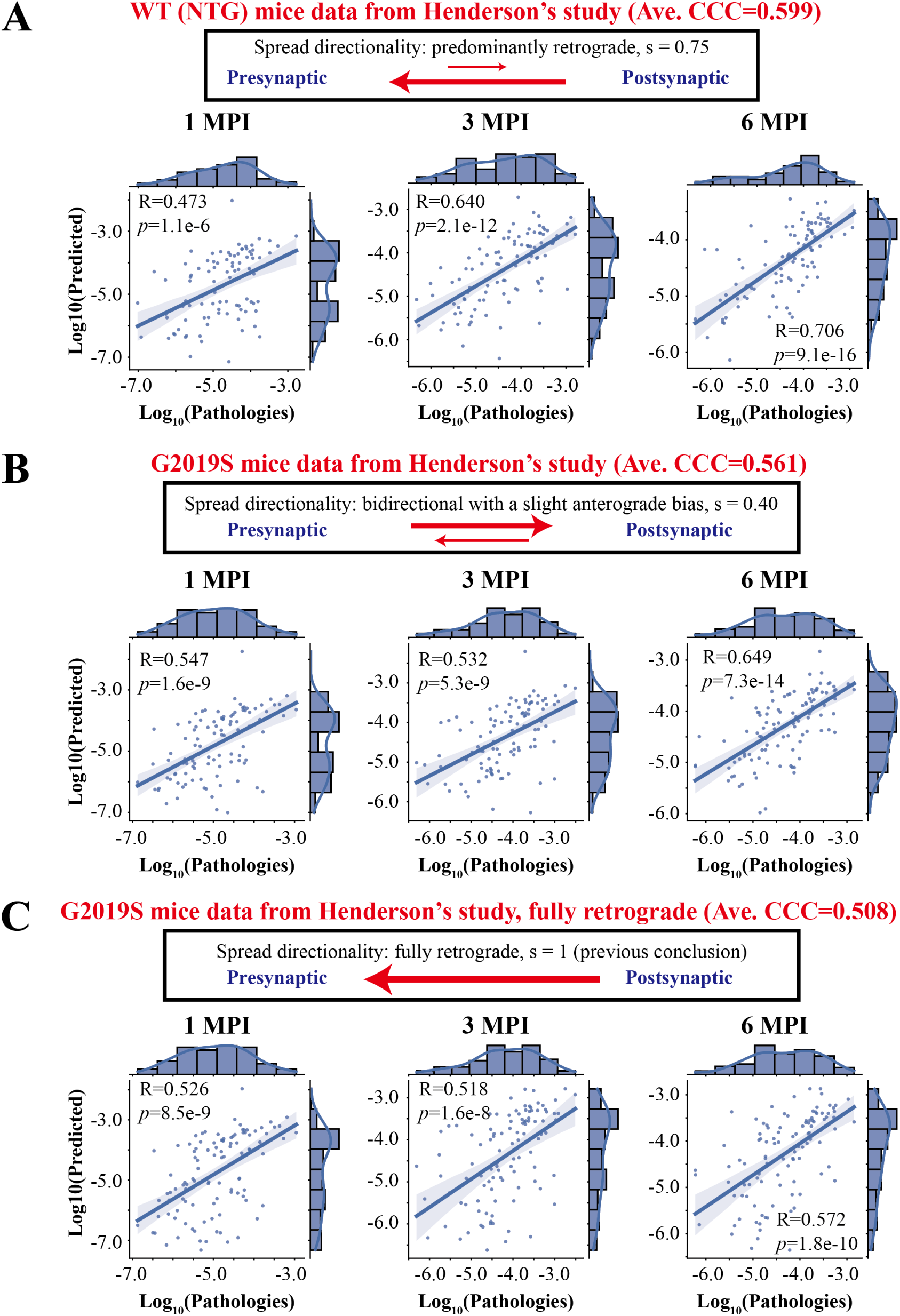
Wild-type non-transgenic (WT) mice and *G2019S LRRK2* (*G2019S*) mice exhibit different directional preferences of pathological α-Syn spread. We used data from Henderson et al. (Henderson et al., 2019a) and our global spread model (see **Methods**). **A**, Best model results for the WT mice (Ave. CCC = 0.599) with the directionality parameter of s=0.75 (predominantly retrograde, in line with Fig. 2A). **B**, Best model results for the *G2019S* mice (Ave. CCC = 0.561) with the directionality parameter of s=0.40 (bidirectional, with a slight anterograde bias). **C**, Model results for the *G2019S* mice (Ave. CCC=0.508) with s=1, which enforces fully retrograde spread as in previous models (Henderson et al., 2019a). The directionality of pathological α-Syn spread is shown by the arrows in the box, while the length of the line indicates the proportion of spread direction. Each dot represents one brain region and the x-axis and y-axis represent the pathology (log10-transformed) found empirically and predicted by the model, respectively. For each situation, the Pearson’s correlation coefficient and the best regression lines for 1, 3 and 6 MPI are also displayed. The shaded ribbon represented the 95% prediction interval. Abbreviations: R, Pearson’s correlation coefficient; *p*, *p* values from linear regression. The detailed description of the datasets was shown in **Methods**.

**Figure S8.**
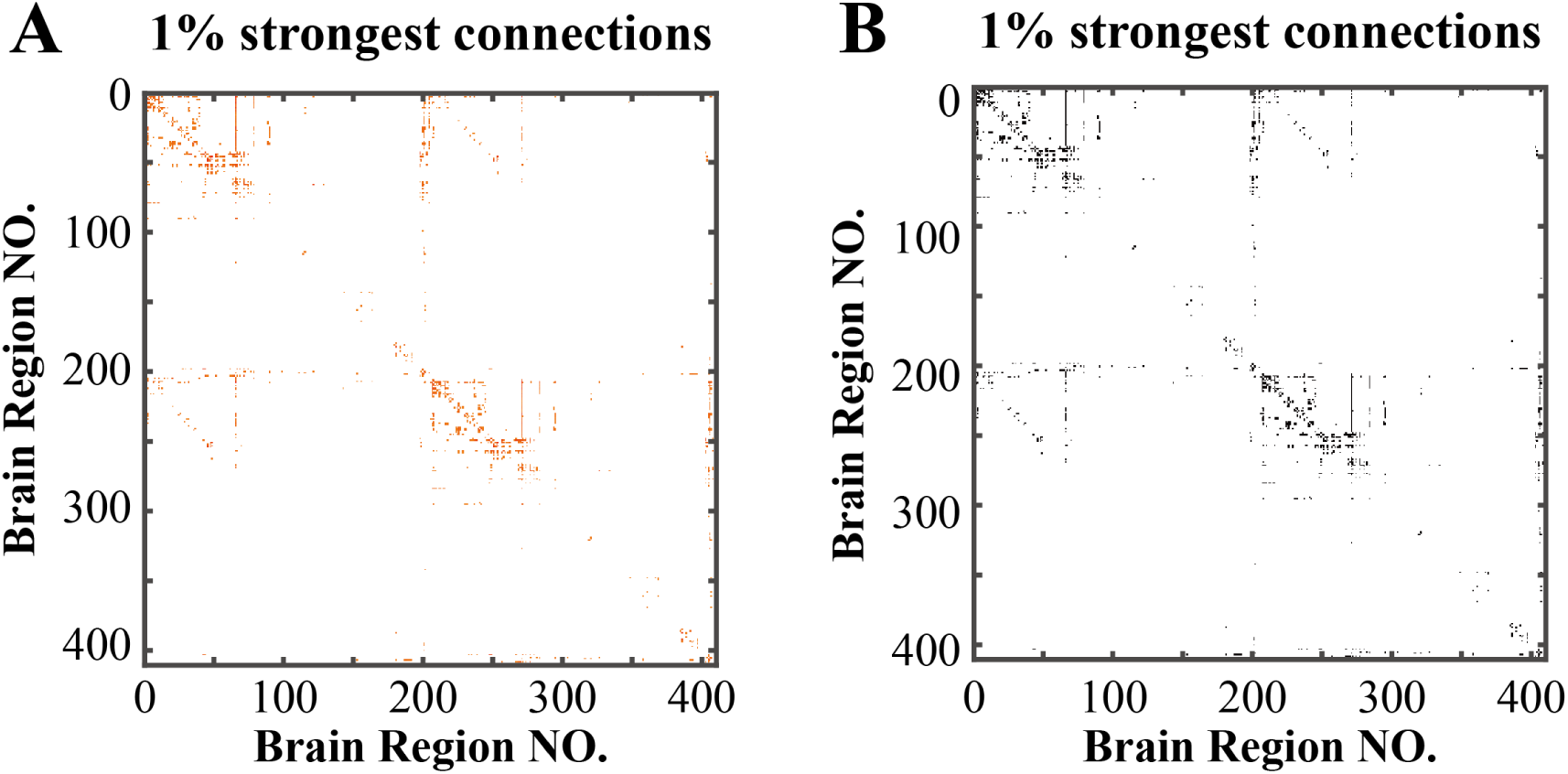
Visualization of connectomes of the 1% of the strongest connections. **A**, Heatmap of this partial connectome, where the warmer colors represent stronger connections (log10(raw_value+1)- transformed). **B**, Heatmap of the adjacency matrix corresponding to **A**.

**Figure S9.**
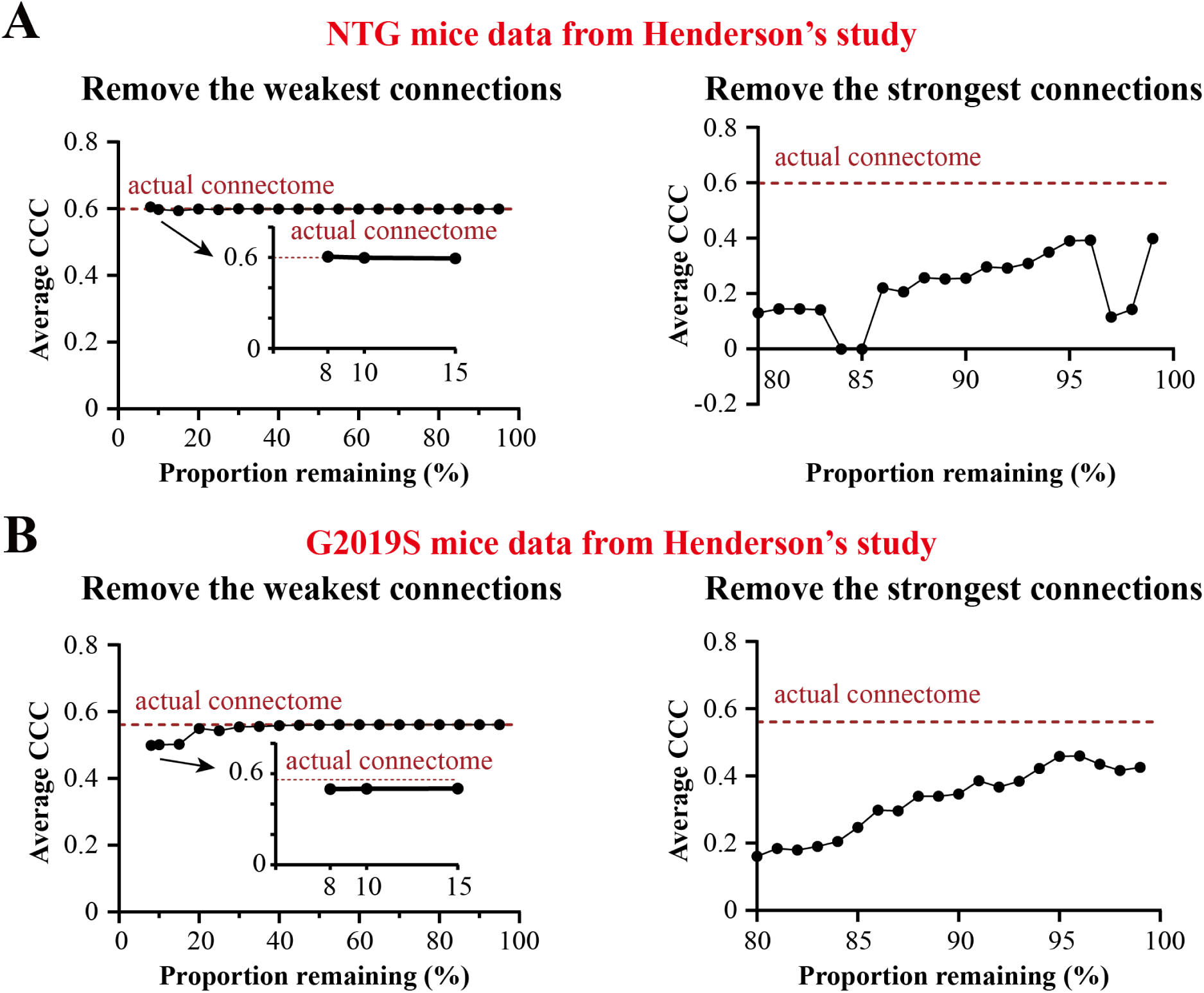
The spread of pathological α-Syn was driven by the partial ‘masked’ connectome composed of the strongest connections for WT mice and *G2019S* mice. The two groups of mice data are from Henderson et al. (Henderson et al., 2019a). **A**, Model performance was not affected when removing up to 92% of the weakest connections (*left*), but was markedly worse after removing the strongest 1% of connections (*right*). The dotted line of the actual connectome showed the best model fitting of the global spread model with the actual ‘masked’ connectome (Ave. CCC = 0.599). **B**, Model fitting results after removing proportions of edges from the connectome in order of connection strength using *G2019S* mouse data from Henderson et al., similar to **A**. The dotted line of the actual connectome showed the best model fitting of the global spread model with the actual ‘masked’ connectome (Ave. CCC = 0.561).

**Figure S10.**
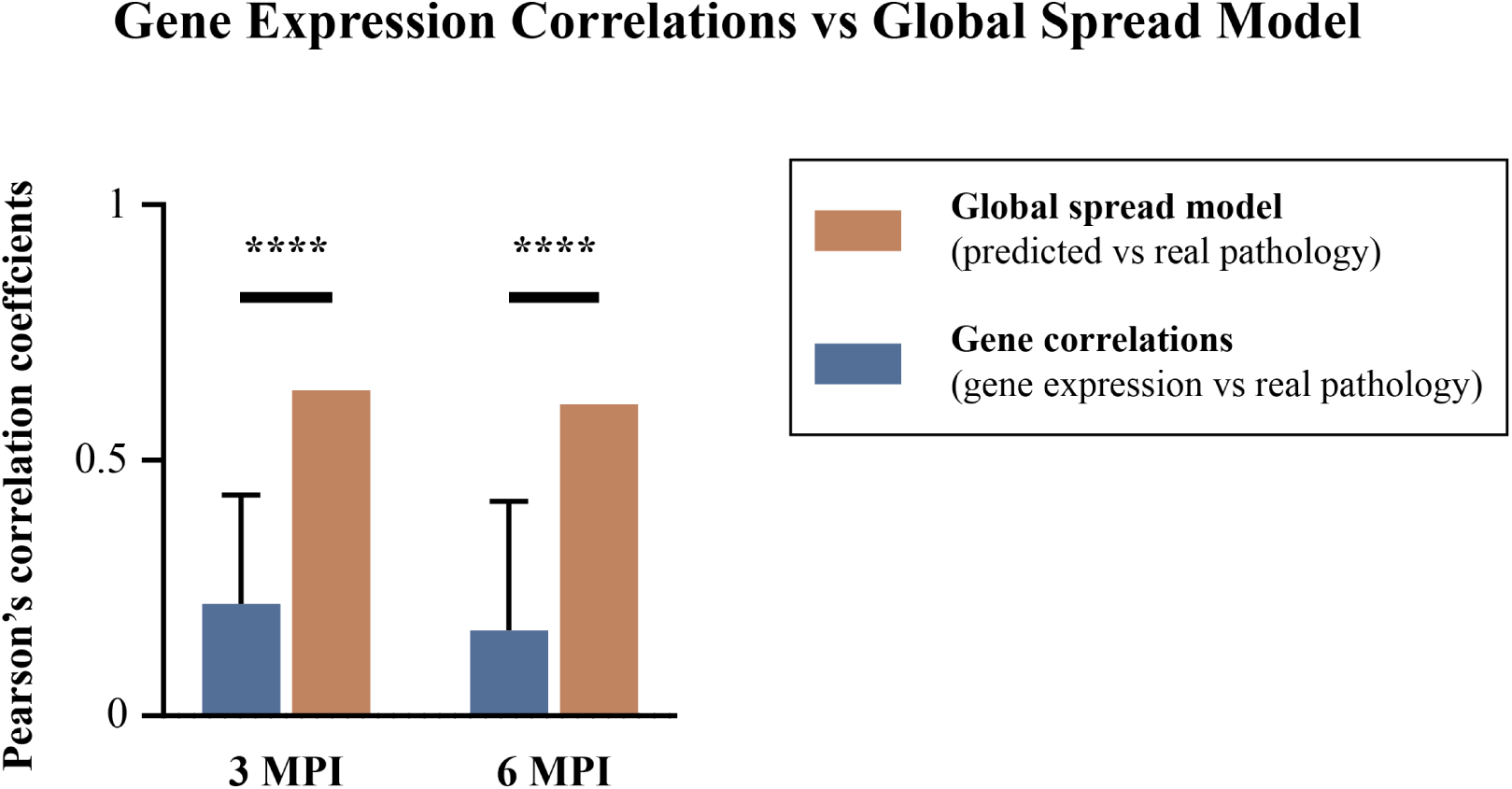
Comparisons of gene expression correlations (defined as the Pearson’s correlation coefficients of gene expression and pathology) and global spread model (the Pearson’s correlation coefficients of model-predicted pathology and real pathology) at 3 and 6 MPI. Gene expression correlations had statistically worse performance than global spread model at both 3 MPI (*p*<0.0001) and 6 MPI (*p*<0.0001) with one sample Wilcoxon test. ****, *p*<0.0001. Data of gene expression correlations were shown as mean±s.d.

**Figure S11.**
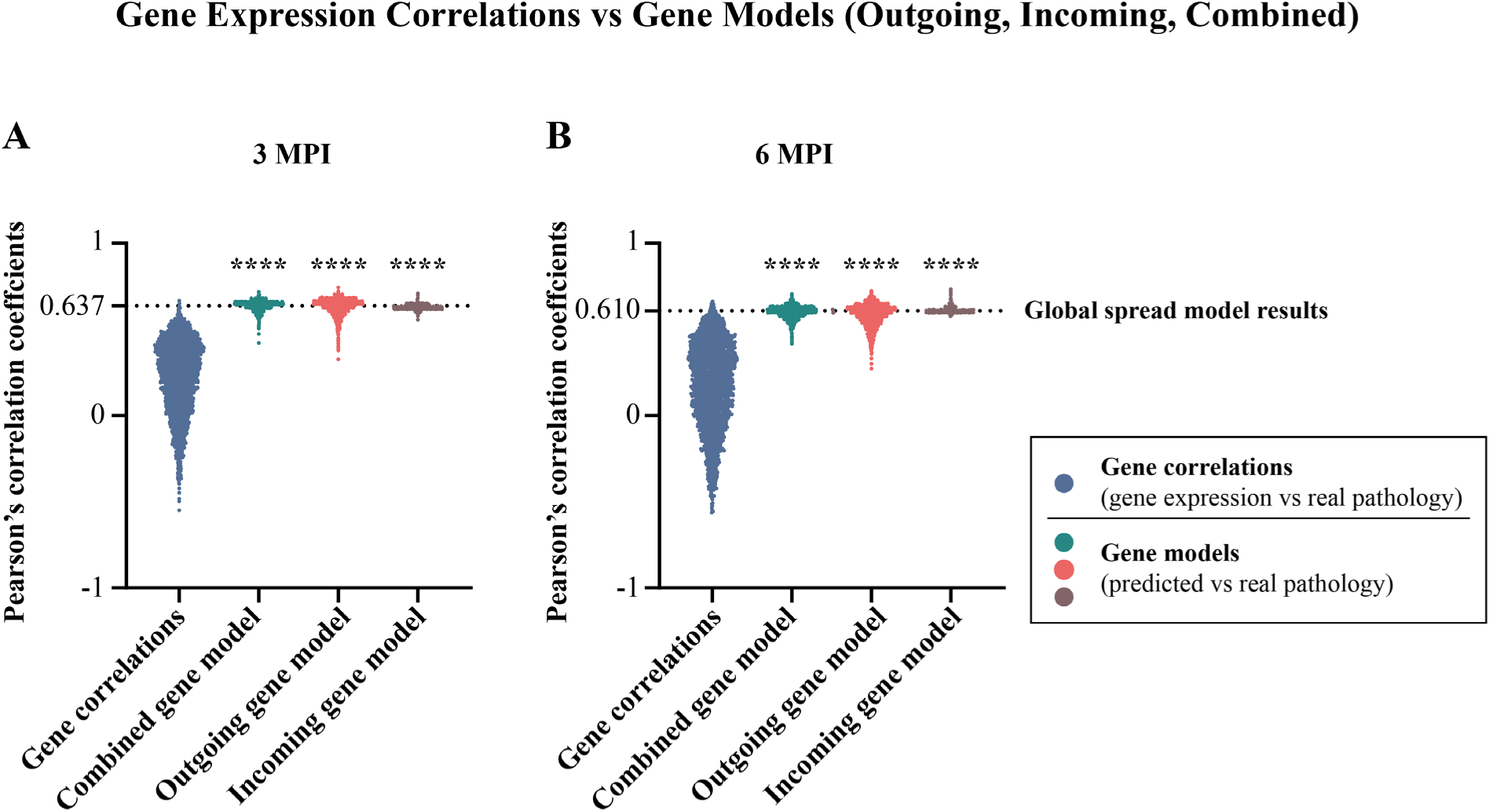
Comparisons of gene expression correlations (defined as the Pearson’s correlation coefficients of gene expression and pathology) and gene models (the Pearson’s correlation coefficients of model-predicted pathology and real pathology; for outgoing, incoming, and combined effects) at 3 (**A**) and 6 (**B**) MPI. Gene expression correlations had statistically worse performance than either of the three gene models at both 3 MPI (*p*<0.0001) and 6 MPI (*p*<0.0001) with Friedman test with Dunn’s multiple comparisons. Each dot in figures represents the result of one individual gene. The dashed lines represent the Pearson’s correlation coefficients of global spread model. ****, *p*<0.0001.

**Figure S12.**
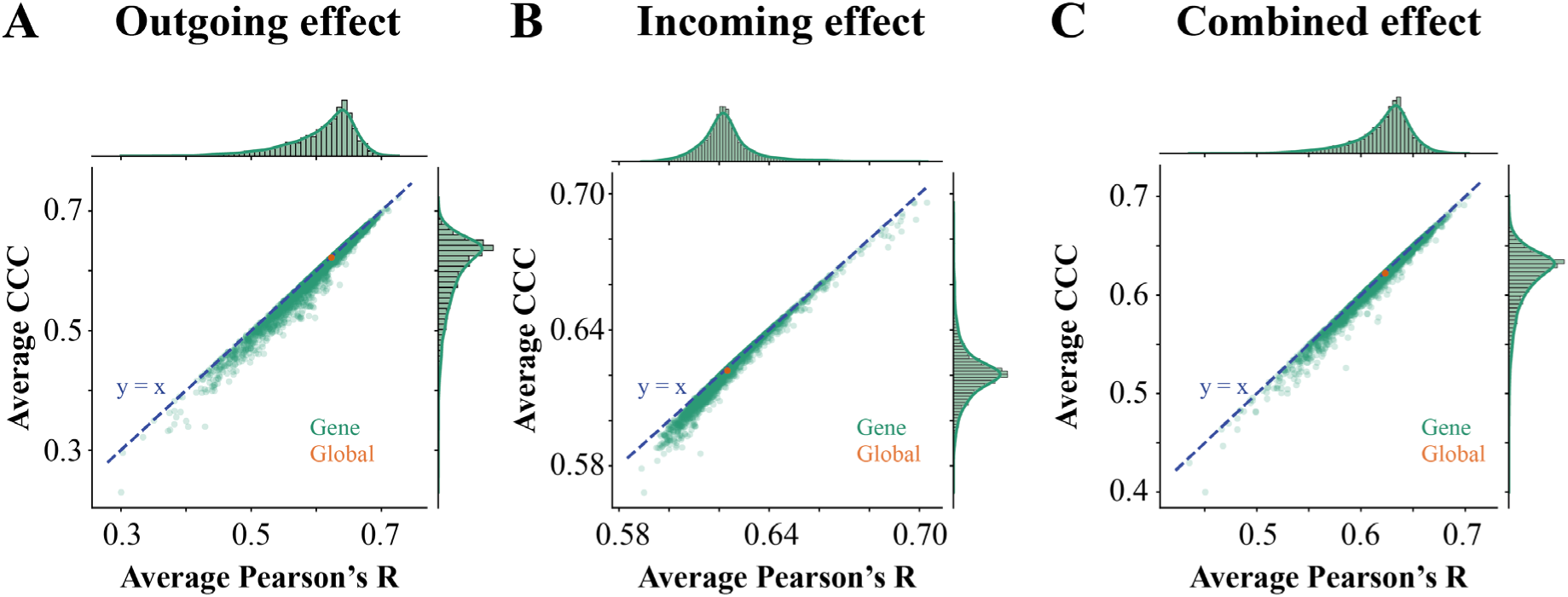
The Ave. CCC vs. the Pearson’s correlation coefficient. Each green dot represents the result of one individual gene and the orange dot represents that of the global spread model. The x-axis and y-axis represent their average of Pearson’s correlation coefficients and Ave. CCC, respectively. Since all the dots were close to the dashed line (y=x), there was not much difference between the Pearson’s correlation coefficient and CCC, revealing the gene model worked well in capturing the scale features. **A**, Outgoing effect. **B**, Incoming effect. **C**, Combined effect.

**Figure S13.**
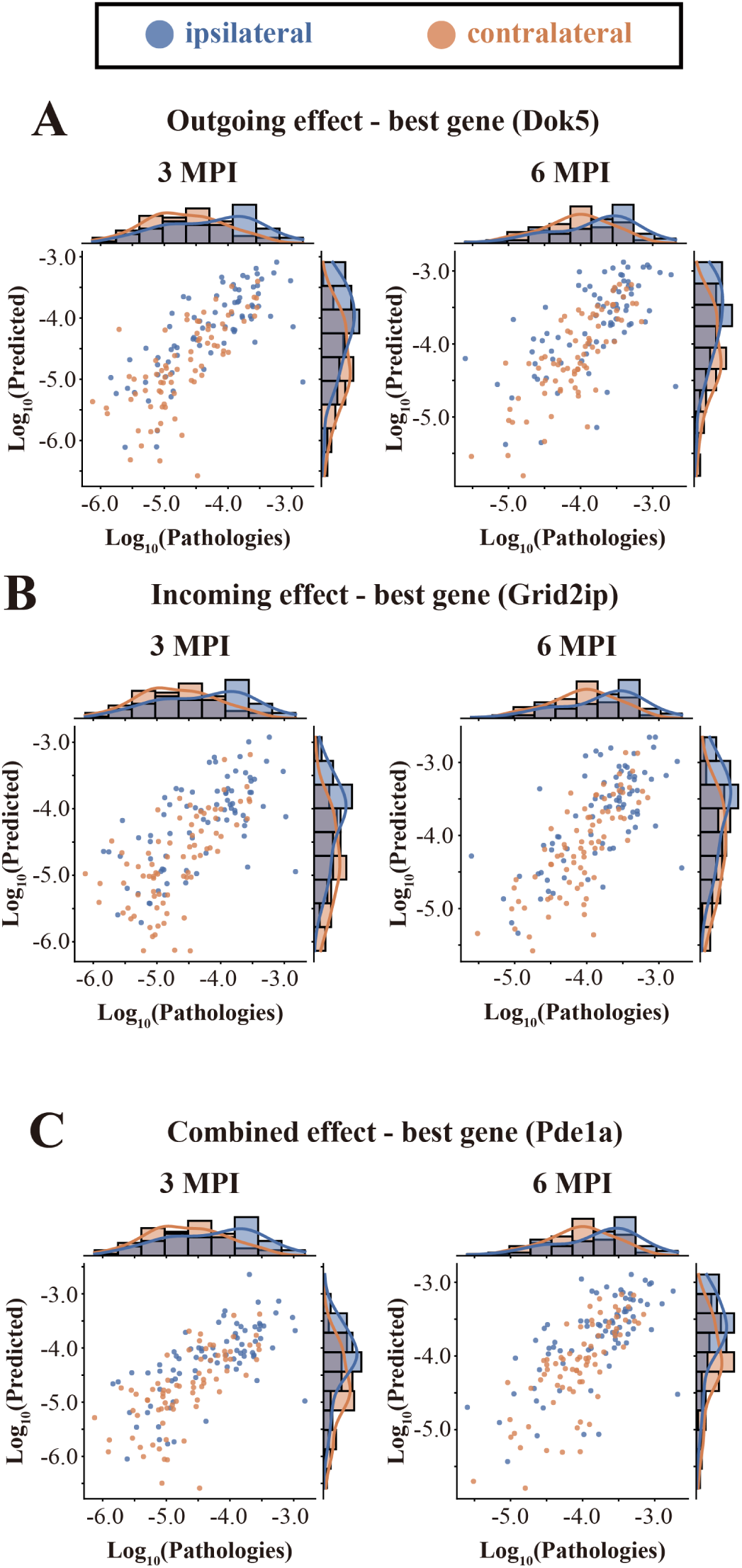
The gene expression model also accurately captures the feature that the ipsilateral hemisphere had more pathology than the contralateral hemisphere both at 3 and 6 MPI. Regions ipsilateral to the injection site are shown in blue, while contralateral regions are shown in orange, with histograms of the univariate distributions shown along the axes. **A**, Best outgoing effect (*Dok5*). **B**, Best incoming effect gene (*Grid2ip*). **C**, Best combined effect gene (*Pde1a*).

**Figure S14.**
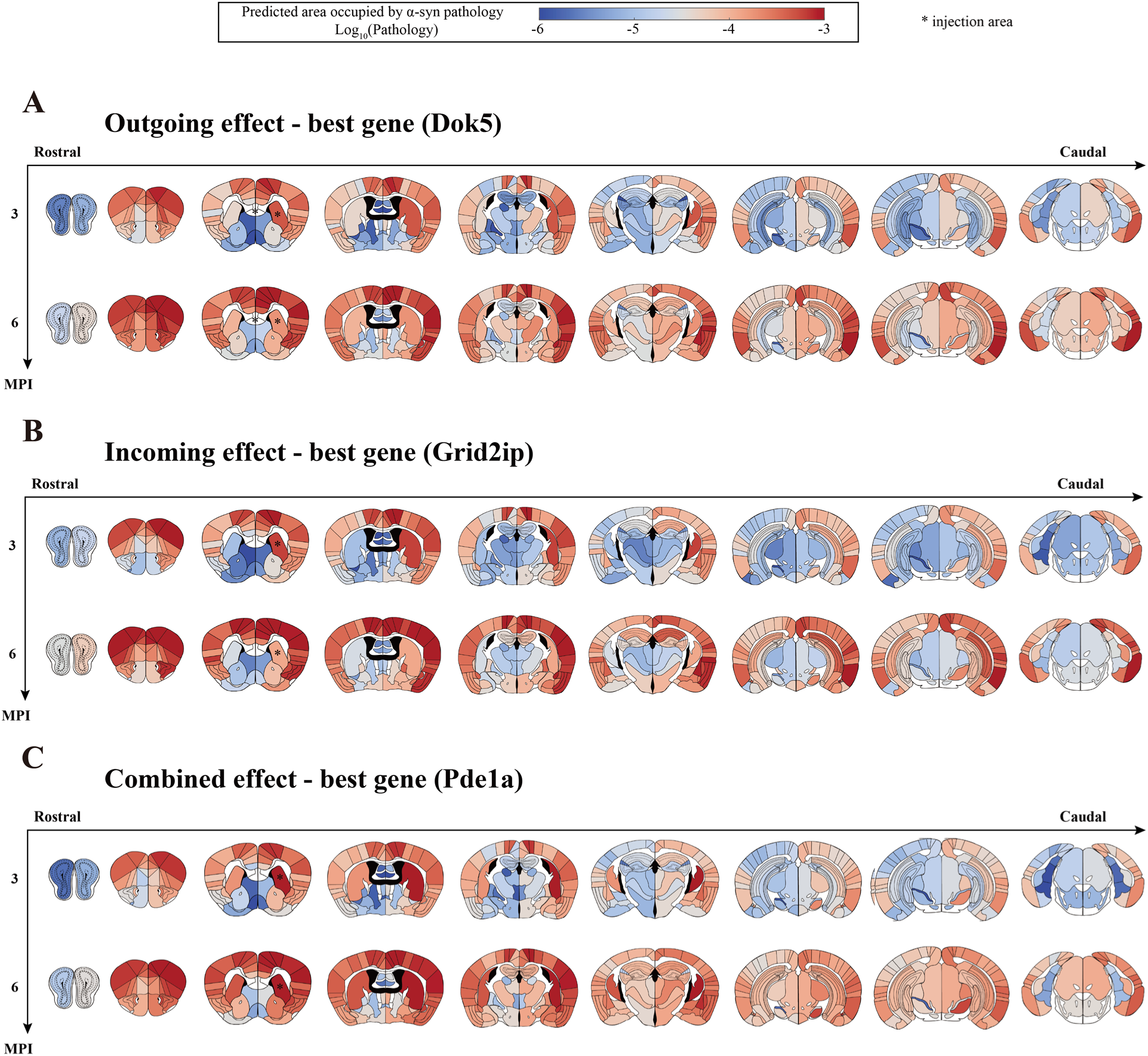
Heatmap of the regional α-Syn pathology predicted by the best gene model (log10- transformed; *, injection area) for **A**, outgoing effect (*Dok5*); **B**, incoming effect (*Grid2ip*). **C**, combined effect (*Pde1a*).

**Figure S15.**
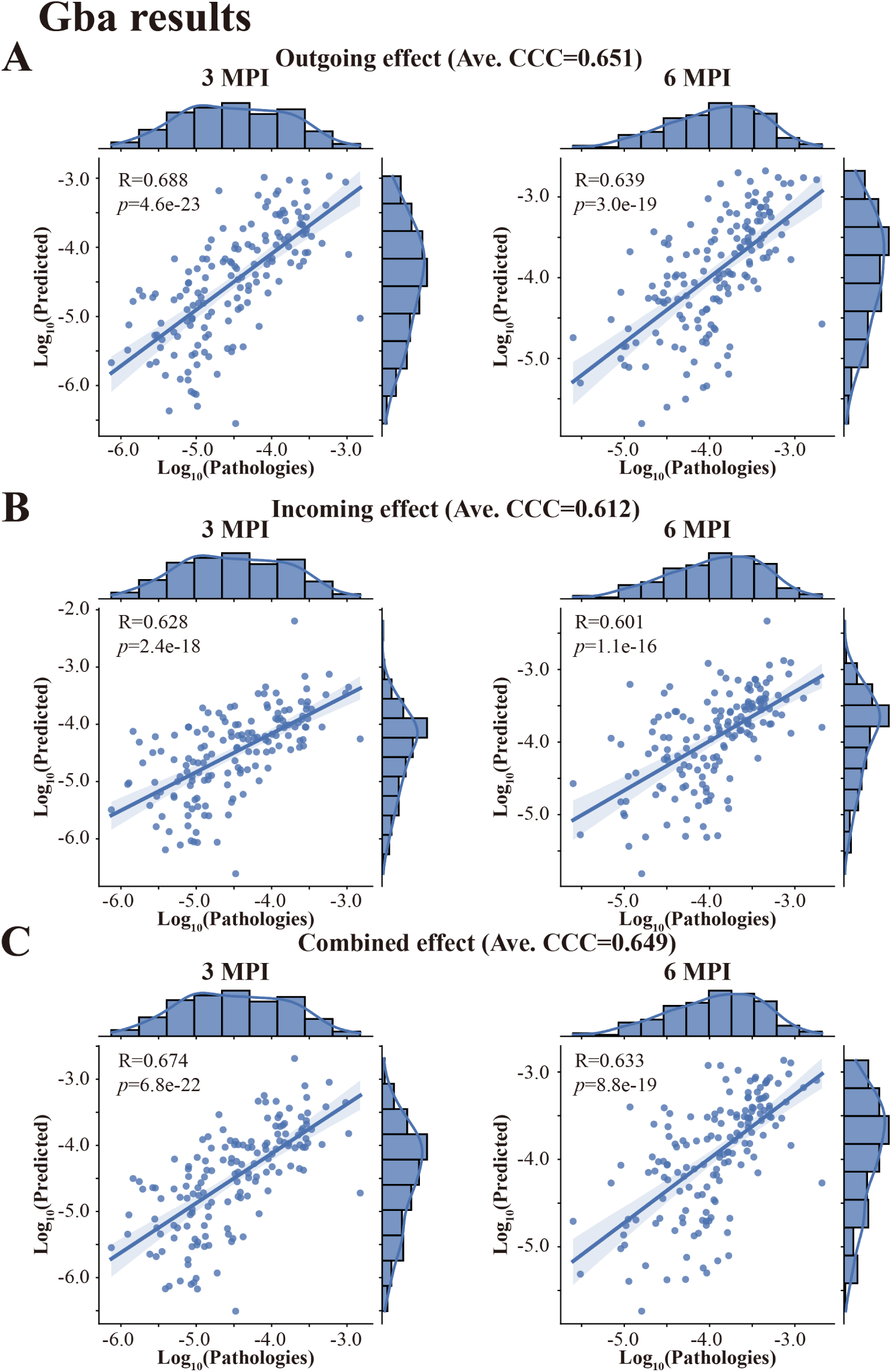
Model results of *Gba* for outgoing (**A**), incoming (**B**), and combined effects (**C**), respectively. Each dot represents one brain region, while the x-axis and y-axis represent the pathology (log10- transformed) found empirically and predicted by the model, respectively. The Pearson’s correlation coefficient and the best regression lines for both 3 and 6 MPI are also displayed. The shaded ribbon represents the 95% prediction interval. Abbreviations: R, Pearson’s correlation coefficient; *p*, *p* values from linear regression.

**Figure S16.**
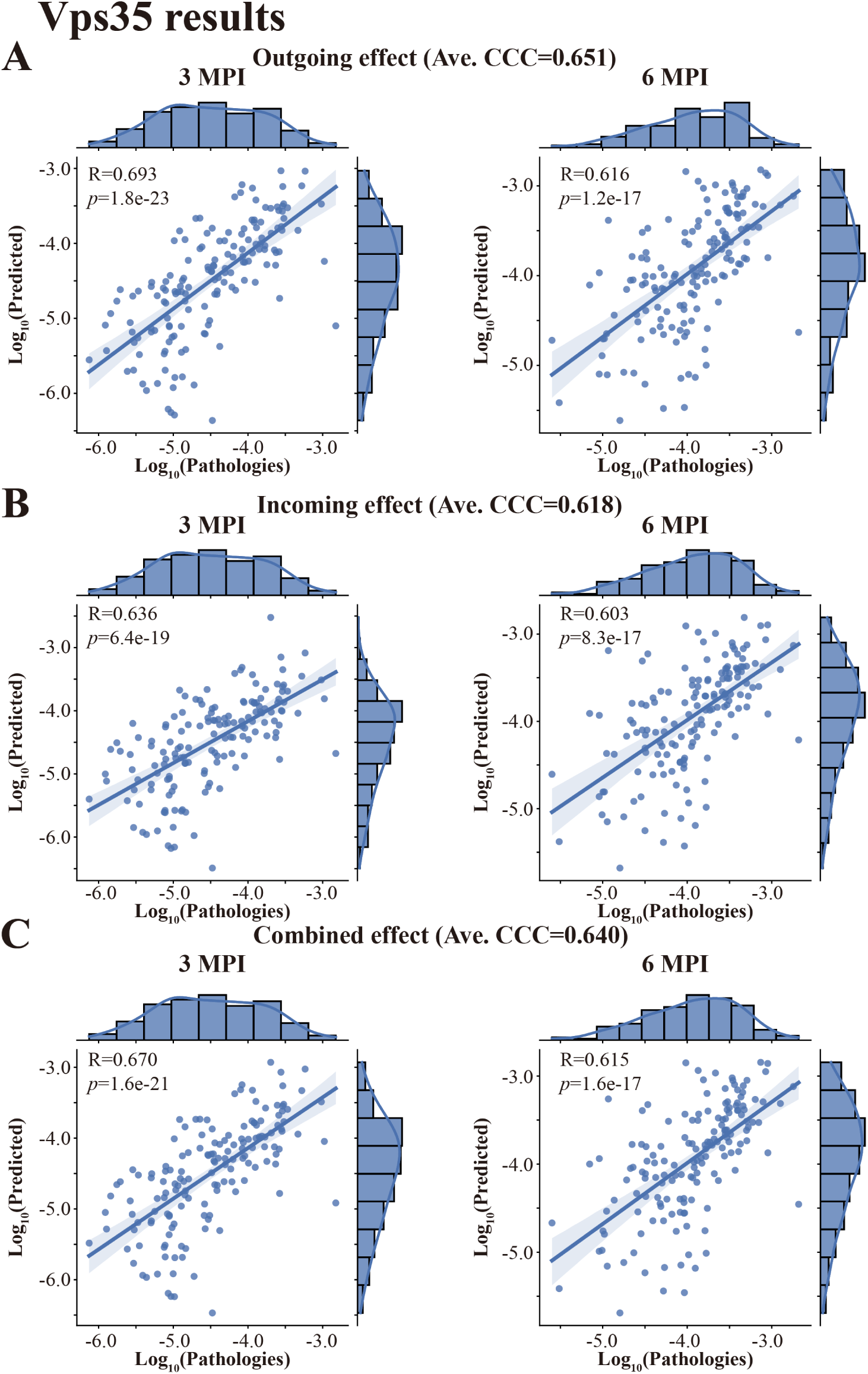
Model results of *Vps35* for outgoing (**A**), incoming (**B**), and combined effect (**C**), respectively. Each dot represents one brain region, while the x-axis and y-axis represent the pathology (log10-transformed) found empirically and predicted by the model, respectively. The Pearson’s correlation coefficient and the best regression lines for both 3 and 6 MPI are also displayed. The shaded ribbon represents the 95% prediction interval. Abbreviations: R, Pearson’s correlation coefficient; *p*, *p* values from linear regression.

**Figure S17.**
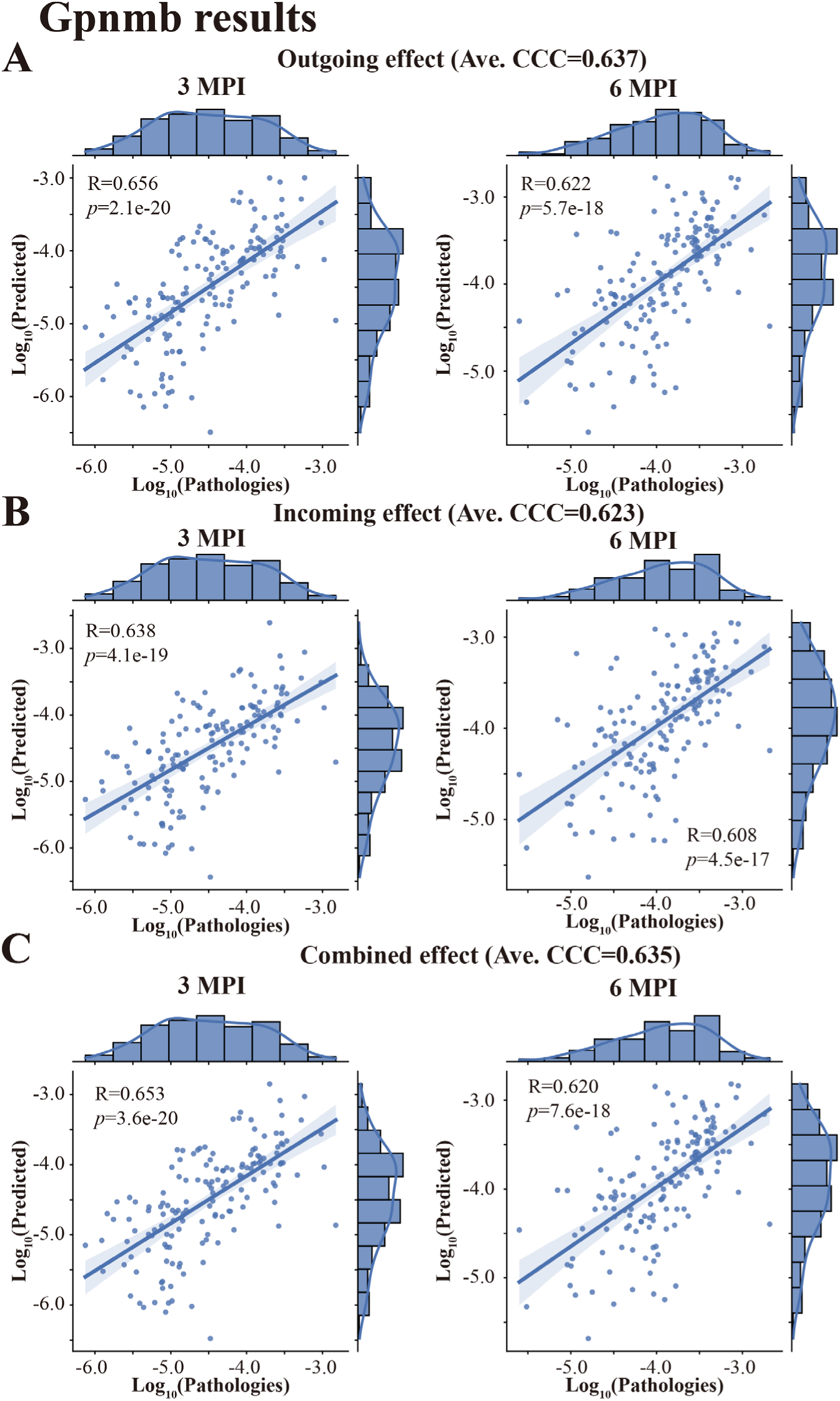
Model results of *Gpnmb* for outgoing (**A**), incoming (**B**), and combined effect (**C**), respectively. Each dot represents one brain region, while the x-axis and y-axis represent the pathology (log10-transformed) found empirically and predicted by the model, respectively. The Pearson’s correlation coefficient and the best regression lines for both 3 and 6 MPI are also displayed. The shaded ribbon represents the 95% prediction interval. Abbreviations: R, Pearson’s correlation coefficient; *p*, *p* values from linear regression.

**Figure S18.**
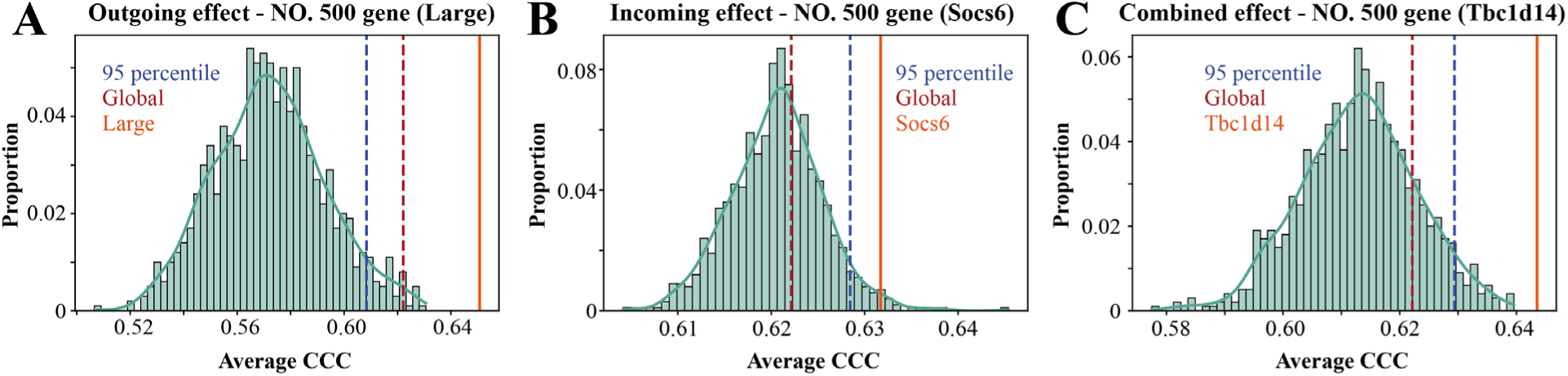
Bootstrapping results with 1,000 replicates in which the elements of the 500th-best genes for each of the **A**, outgoing (*Large*); **B**, incoming (*Socs6*). **C**, combined (*Tbc1d14*) effects, respectively, were randomly permuted. The true Ave. CCC of 500th-best genes were all higher than the 95% percentiles of the null model distributions.

**Figure S19.**
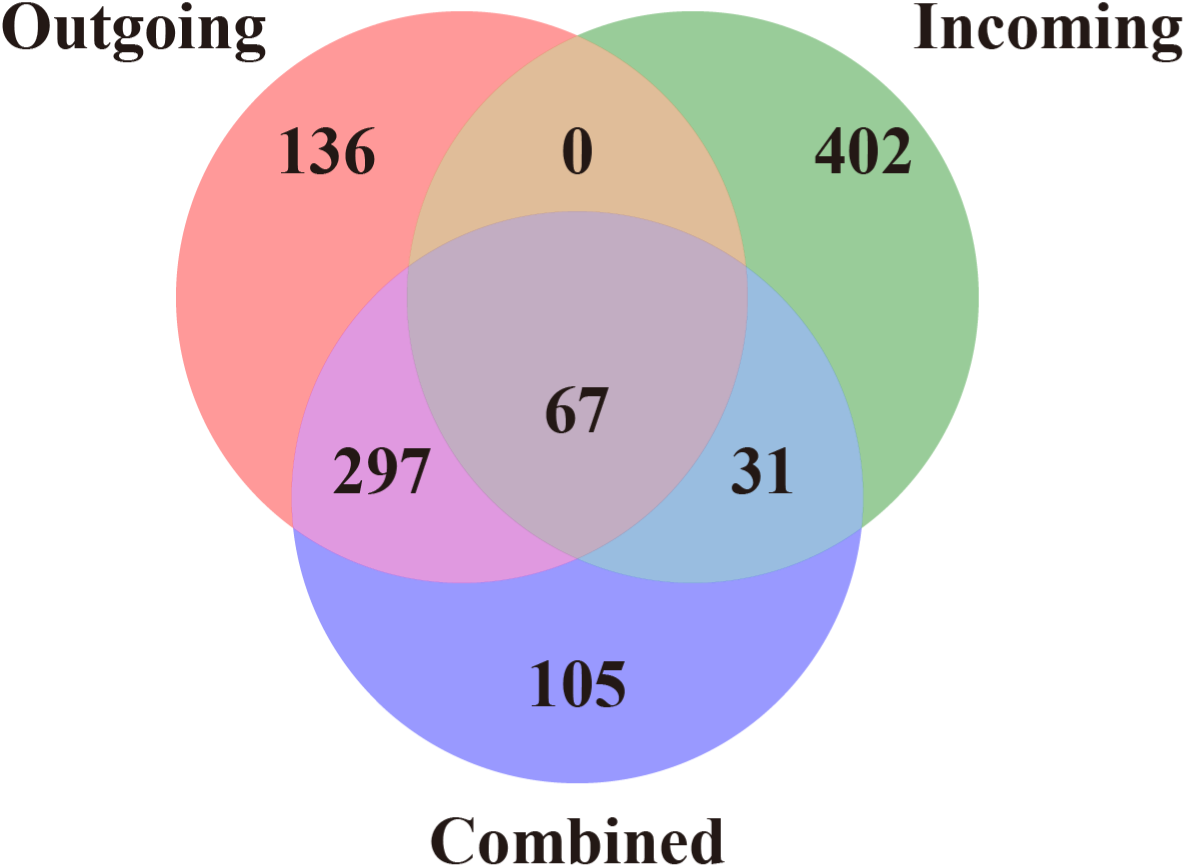
Venn diagram showing the relationships among the top 500 genes of the outgoing, incoming, and combined effects. There was more gene overlap between the outgoing effect and the combined effect than the other effect pairs.

**Figure S20.**
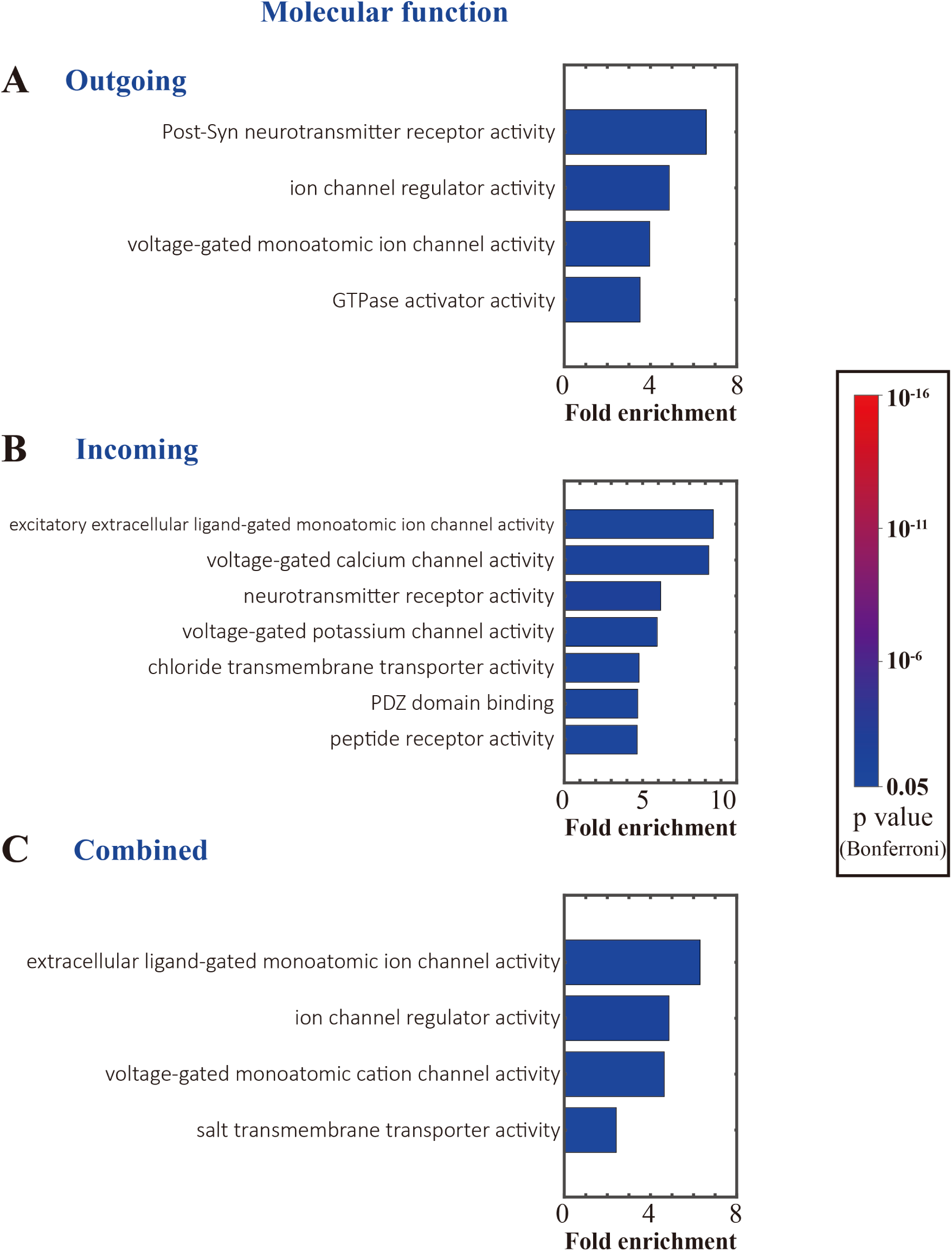
Molecular function of GO analyses for the top 500 genes of outgoing, incoming, and combined effect. **A**, Outgoing effect. **B**, Incoming effect. **C**, Combined effect. For each GO entry, the displayed results were strictly sorted by the highest fold enrichment and statistically significant after Bonferroni correction. The colors of the bars represent the *p* values, and the length represents the fold enrichment. Only the primary hierarchy is shown. Abbreviations: Post-Syn, postsynaptic.

**Figure S21.**
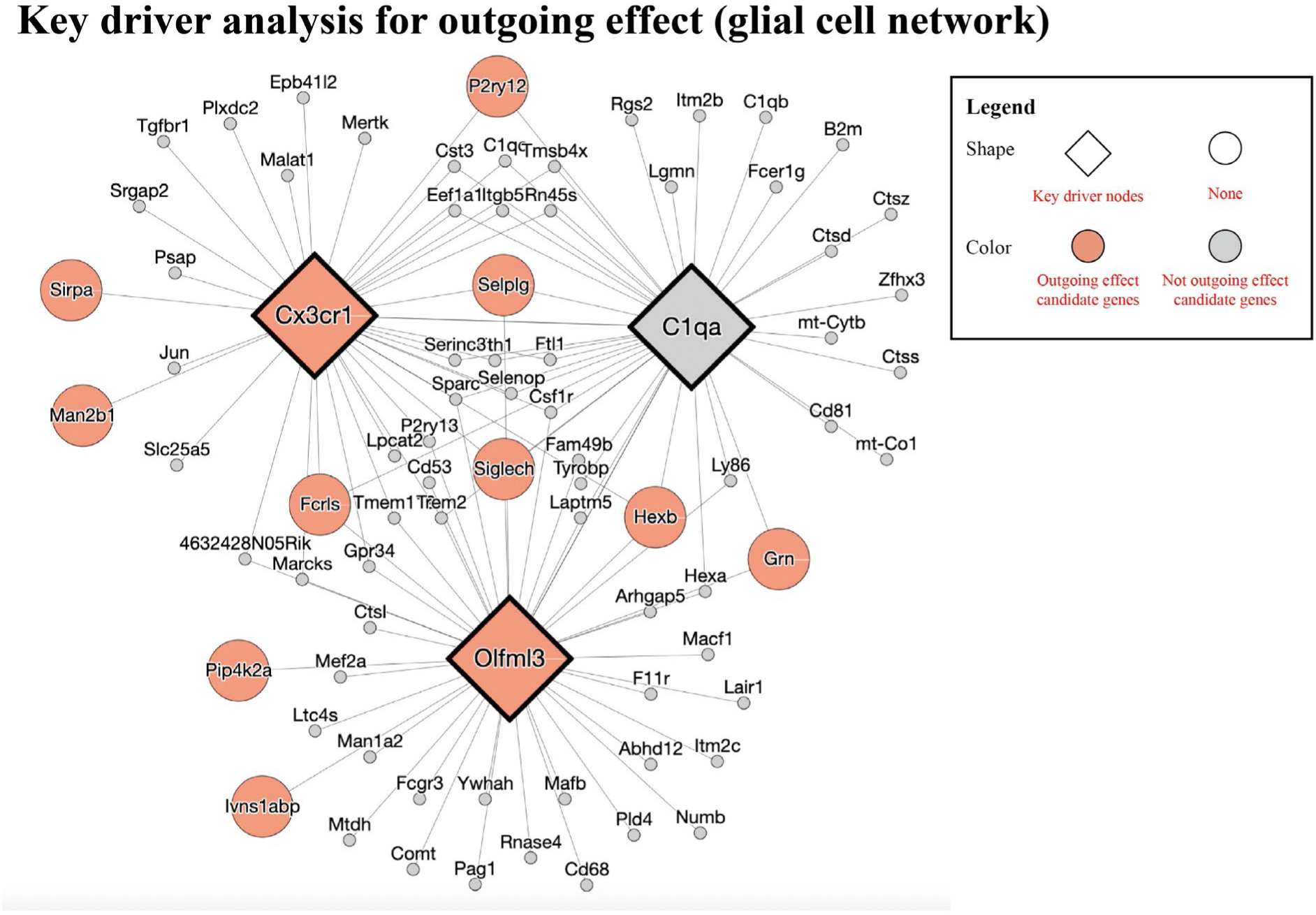
KD analysis for candidate genes of incoming effect and the associated gene regulatory network on glial cell scRNAseq data. The diamond shape indicate genes that were identified to be KDs, and circles indicate not. The colors indicate whether the gene occurs in outgoing effect candidate gene subset or not. The SCING neuronal network is constructed using scRNAseq data from Tabula Muris, Tabula Muris Senis, and Mouse Cell Atlas, with networks built on the Allen 10X dataset from the hippocampus, visual cortex, somatosensory cortex, and primary motor cortex brain regions.

**Figure S22.**
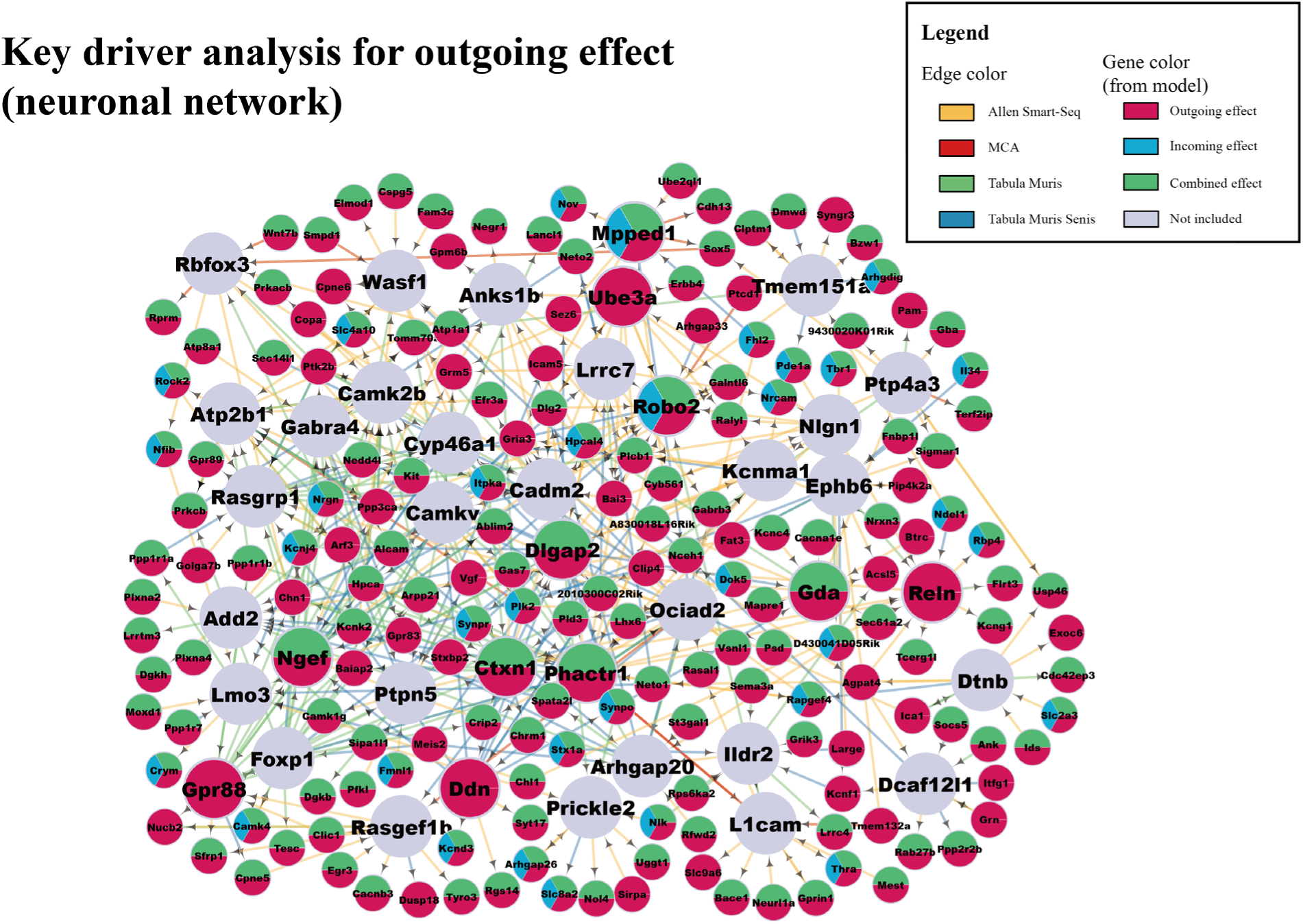
KD analysis for candidate genes of outgoing effect and the associated gene regulatory network on neuronal scRNAseq data. The larger circles indicate genes that were identified to be KDs, and smaller circles indicate candidate genes from the outgoing effects. The direction of edges indicates the regulatory relationship between genes in the SCING neuronal network constructed using scRNAseq data from single cell atlases Allen Smart-seq, mouse cell atlas (MCA), Tabula Muris, and Tabula Muris Senis. The color inside the circle denotes whether the gene has outgoing (red), incoming (green), or combined (blue) effect, while the multiple colors indicate the gene is from multiple categories. KD genes that did not appear in any of the effects were marked as gray color. Note that network KDs for the candidate genes do not have to be candidate genes themselves but are predicted to regulate the candidate genes.

**Figure S23.**
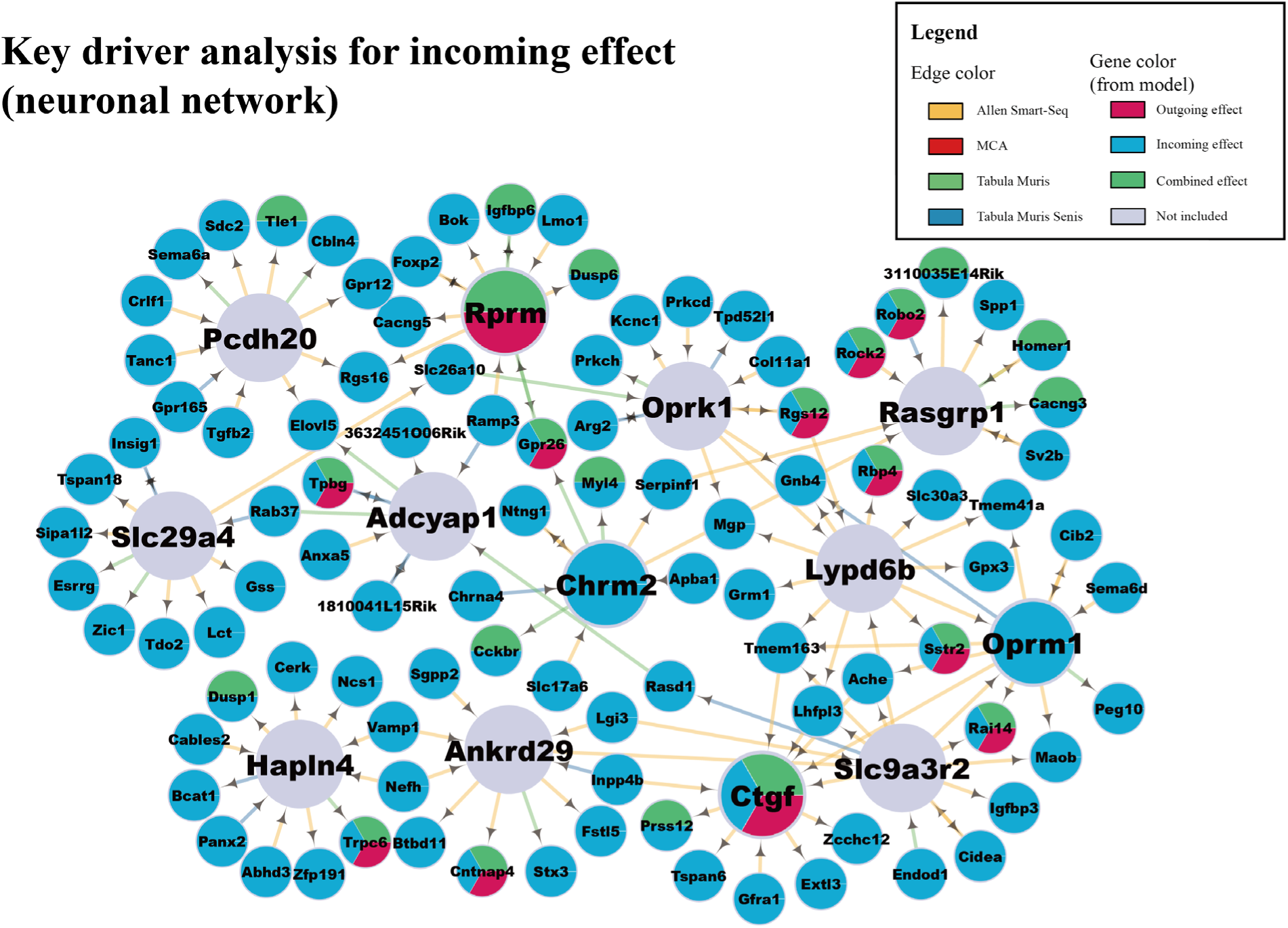
KD analysis for candidate genes of incoming effect and the associated gene regulatory network on neuronal scRNAseq data. The larger circles indicate genes that were identified to be KDs, and smaller circles indicate candidate genes from the incoming effects. The direction of edges indicates the regulatory relationship between genes in the SCING neuronal network constructed using scRNAseq data from single cell atlases Allen Smart-seq, mouse cell atlas (MCA), Tabula Muris, and Tabula Muris Senis. The color inside the circle denotes whether the gene has outgoing (red), incoming (green), or combined (blue) effect, while the multiple colors indicate the gene is from multiple categories. KD genes that did not appear in any of the effects were marked as gray color. Note that network KDs for the candidate genes do not have to be candidate genes themselves but are predicted to regulate the candidate genes.

**Figure S24.**
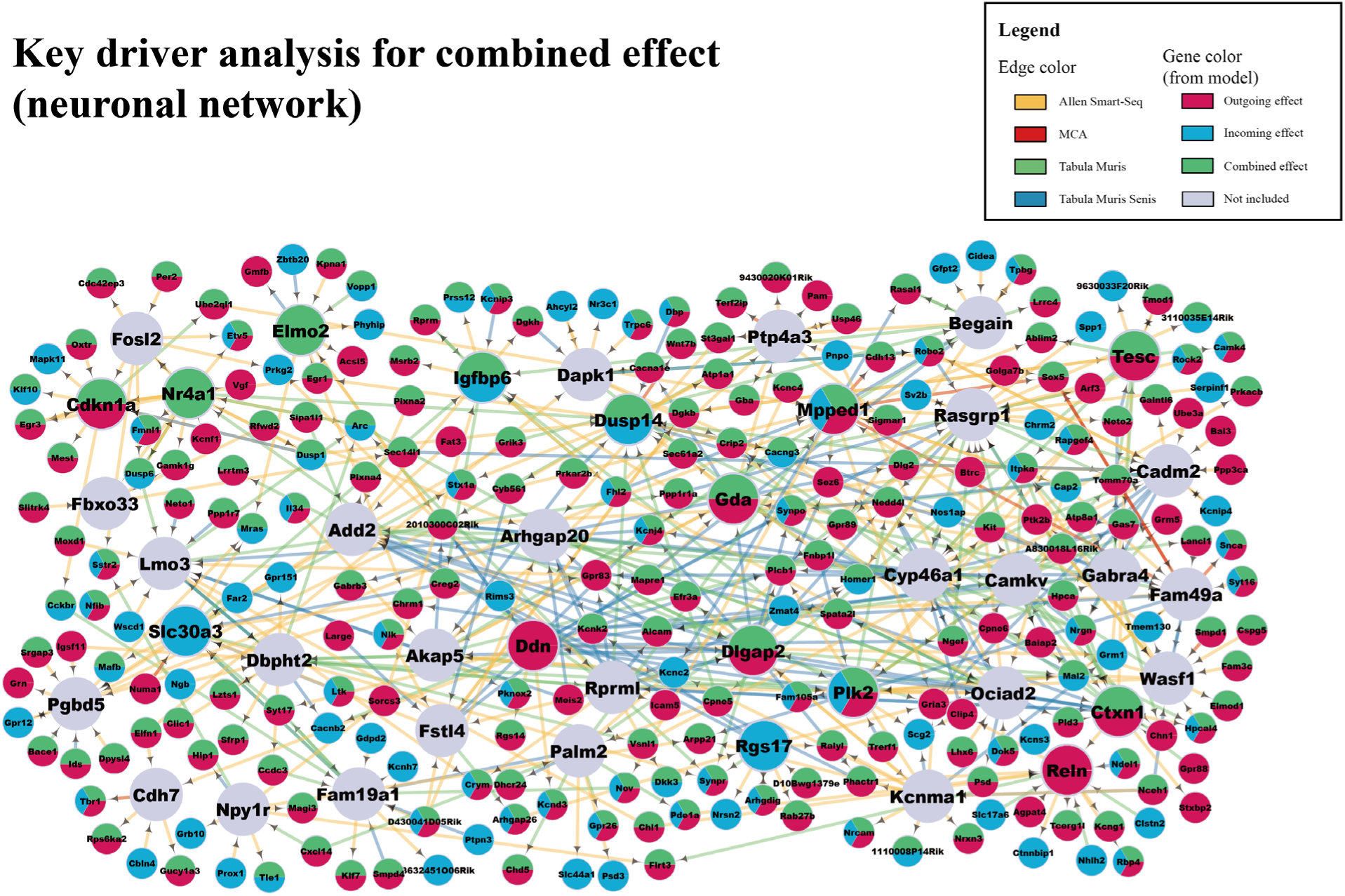
KD analysis for candidate genes of combined effect and the associated gene regulatory network on neuronal scRNAseq data. The larger circles indicate genes that were identified to be KDs, and smaller circles indicate candidate genes from the combined effects. The direction of edges indicates the regulatory relationship between genes in the SCING neuronal network constructed using scRNAseq data from single cell atlases Allen Smart-seq, mouse cell atlas (MCA), Tabula Muris, and Tabula Muris Senis. The color inside the circle denotes whether the gene has outgoing (red), incoming (green), or combined (blue) effect, while the multiple colors indicate the gene is from multiple categories. KD genes that did not appear in any of the effects were marked as gray color. Note that network KDs for the candidate genes do not have to be candidate genes themselves but are predicted to regulate the candidate genes.

**Figure S25.**
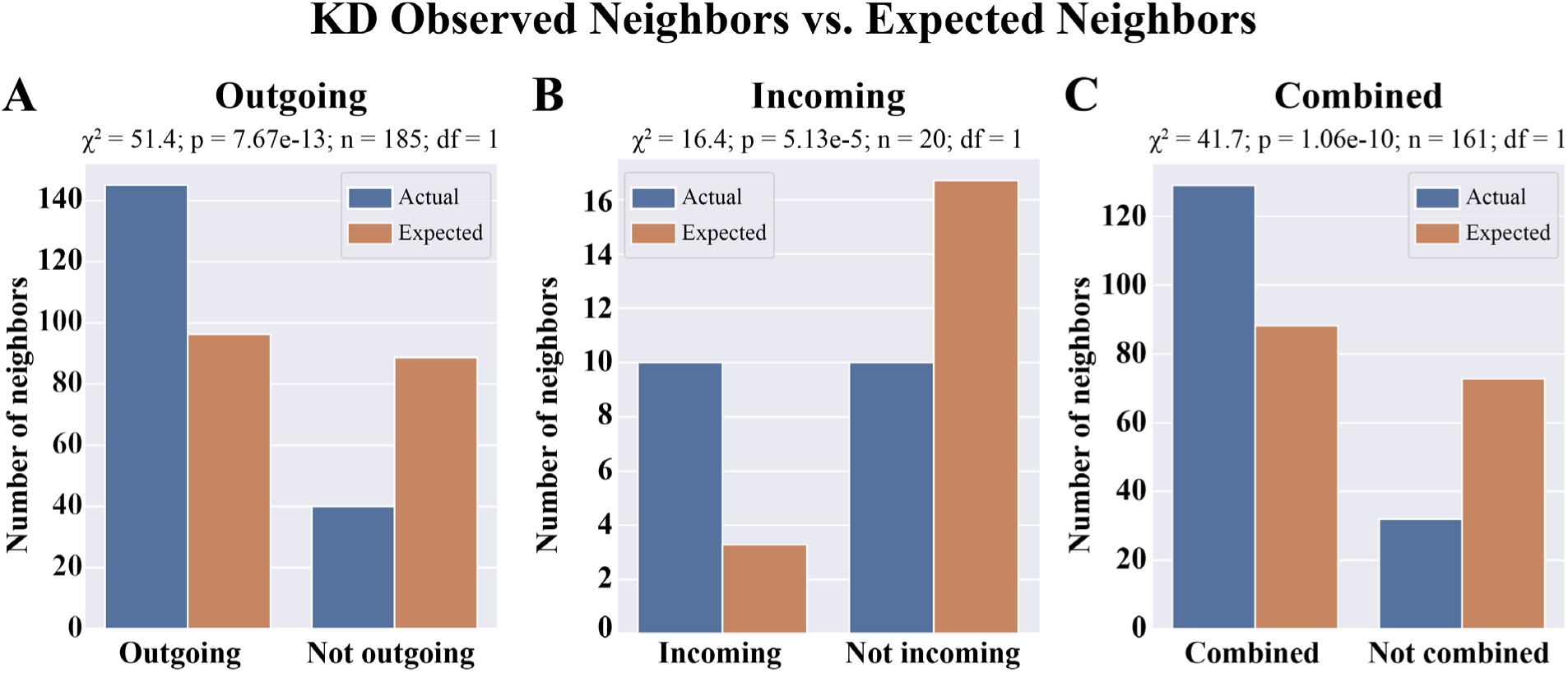
Chi-square goodness of fit test for number of neighbors of each category for the union regulatory network. A, Outgoing subset. B, Incoming subset. C, Combined subset.

